# The elusive neural signature of emotion regulation capabilities: evidence from a large-scale consortium

**DOI:** 10.1101/2025.08.18.670843

**Authors:** Maurizio Sicorello, Jenny Zaehringer, Lena Paschke, Rosa Steimke, Christine Stelzel, Peter J. Gianaros, Kevin S. LaBar, John L. Graner, Sang H. Kim, Michèle Wessa, Magdalena Sandner, Franziska Weinmar, Birgit Derntl, Thomas E. Kraynak, Nathan T.M. Huneke, Harry Fagan, Nils Kohn, Guillén Fernández, Linlin Yan, Agar Marín-Morales, Juan Verdejo-Román, Trevor Steward, Ben J. Harrison, Christopher G. Davey, Denise Dörfel, Henrik Walter, Maital Neta, Jordan Pierce, David S. Stolz, Johanna Kissler, Anissa Benzait, Susanne Erk, Stella Berboth, Carien M. van Reekum, Emma Tupitsa, Satja Mulej Bratec, Christian Sorg, Laura Müller-Pinzler, Andrzej Sokołowski, Wojciech Ł. Dragan, Monika Folkierska-Żukowska, Valerie L. Jentsch, Christian J. Merz, Christoph Scheffel, Kersten Diers, Kaoru Nashiro, Steve Heinke, Jungwon Min, Mara Mather, Anne Gärtner, Kateri McRae, John P. Powers, Nick Doren, Silvia U. Maier, Stephan Nebe, Isabel Dziobek, Michael Gaebler, Judith K. Daniels, Matthias Burghart, Stephanie N. L. Schmidt, Lena Hofhansel, Ute Habel, Carmen Morawetz

**Affiliations:** Department of Psychosomatic Medicine and Psychotherapy, Central Institute of Mental Health, Medical Faculty Mannheim, Heidelberg University, Germany; German Center for Mental Health (DZPG), Partner Site Mannheim-Heidelberg-Ulm; Charité - Universitätsmedizin Berlin, Department of Psychiatry and Neurosciences, Division Mind and Brain, Berlin, Germany; Department of Psychology, Division of Clinical Psychology and Psychotherapy in Adults, University of Bremen, Bremen, Germany; International Psychonanalytic University Berlin, Berlin, Germany; University of Pittsburgh, Department of Psychology, Center for Mind-Body Science and Health; Center for Cognitive Neuroscience, Duke University, Durham, NC USA; Department of Brain and Cognitive Engineering, Korea University; Central Institute of Mental Health, Department of Neuropsychology and Psychological Resilience Research, Mannheim, Germany; DKFZ Hector Cancer Institute at the University Medical Center Mannheim, Germany; German Cancer Research Center (DKFZ) Heidelberg, Division Cancer Survivorship and Psychological Resilience, Germany; Johannes Gutenberg-University Mainz, Department of Clinical Psychology and Neuropsychology, Mainz, Germany; Department of Psychiatry and Psychotherapy, Women’s Mental Health and Brain Function, Tübingen Center for Mental Health (TüCMH), University of Tübingen, Tübingen, Germany.; German Center for Mental Health (DZPG), partner site Tübingen, Germany; Department of Psychiatry, University of Pittsburgh; University Department of Psychiatry, Faculty of Medicine, University of Southampton, UK; Hampshire and Isle of Wight Healthcare National Health Service Foundation Trust, Southampton, UK; Radboud University Medical Center, Department of Medical Neuroscience, Donders Institute for Brain, Cognition and Behaviour, Nijmegen, Netherlands; Department of Social, Developmental and Educational Psychology, University of Huelva, Spain; Department of Personality, Evaluation and Psychological Treatment; Mind Brain and Behavior Research Center (CIMCYC-UGR), University of Granada; The University of Melbourne, Victoria, Australia; TUD Dresden University of Technology, Center for Interdisciplinary Digital Sciences, Dresden, Germany; TUD Dresden University of Technology, Faculty of Psychology, Chair of Differential and Personality Psychology, Dresden, Germany; Department of Psychology, University of Nebraska-Lincoln; Center for Brain, Biology, and Behavior, University of Nebraska-Lincoln; University of Lübeck, Department of Psychiatry and Psychotherapy, Social Neuroscience Lab; Bielefeld University, Department of Psychology, Bielefeld, Germany; Charité Universitätsmedizin Berlin, Department of Psychiatry and Neurosciences, Berlin, Germany; Centre for Integrative Neuroscience and Neurodynamics, School of Psychology and Clinical Language Sciences, University of Reading, Reading, UK; Department of Psychology, Faculty of Arts, University of Maribor, Slovenia; Department of Neuroradiology, Technische Universität München, Munich, Germany; Department of Psychiatry, Technische Universität München, Munich, Germany; TUM-Neuroimaging Center, Technische Universität München, Munich, Germany; Stanford University, Stanford, USA; Department of Psychology, Jagiellonian University, Kraków, Poland; University of Toronto Mississauga, Mississauga, Canada; Ruhr University Bochum, Institute of Cognitive Neuroscience, Department of Cognitive Psychology; Leonard Davis School of Gerontology, University of Southern California, Los Angeles, USA; Human-IST Institute, University of Fribourg, Fribourg, Switzerland; Center of Economic Psychology, University of Basel, Basel, Switzerland; University of Denver, Department of Psychology; University of North Carolina at Chapel Hill, North Carolina Translational and Clinical Sciences Institute; Zurich Center for Neuroeconomics, Department of Economics, University of Zurich, Zurich, Switzerland; Clinical Psychology of Social Interaction, Humboldt-Universität zu Berlin; German Center of Mental Health (DZPG), partner site Berlin-Potsdam, Germany; Max Planck Institute for Human Cognitive and Brain Sciences, Neurology Department, Leipzig, Germany; Charité - Universitätsmedizin Berlin; Department of Clinical Psychology, University of Groningen, The Netherlands; Traumacentrum Beilen, GGZ Drenthe, The Netherlands; University of Konstanz, Department of Clinical Psychology and Psychotherapy, Konstanz, Germany; Max Planck Institute for the Study of Crime, Security and Law, Freiburg im Breisgau, Germany; Department of Psychiatry, Psychotherapy, and Psychosomatics, RWTH Aachen University; Department of Psychology, University of Innsbruck, Austria

## Abstract

Cognitive reappraisal is a fundamental emotion regulation strategy for mental and physical well-being, but how its neural mechanisms relate to individual differences remains poorly understood. In a consortium effort analyzing 40 fMRI datasets (*N*=2,175), we examined the relationship between neural activation during reappraisal tasks and three core individual difference indices of reappraisal capabilities: (1) trait questionnaires, (2) task-based affective ratings, and (3) amygdala down-regulation. Strikingly, there was no shared overlap across these three common indices. Only a very weak correlation emerged between amygdala down-regulation and task-based affective ratings. Whole-brain analyses revealed no reliable neural associations with trait questionnaires, and associations with task-based affective ratings fell outside canonical emotion regulation networks (e.g., prefrontal circuitry). Moreover, amygdala down-regulation, often interpreted as a stable individual marker, was confounded by person-specific whole-brain responses — a limitation extending to fMRI research beyond the emotion regulation domain. These findings challenge the assumption that an individual’s prefrontal activity is a valid indicator of their reappraisal capabilities and suggest that common trait, behavioral, and neural measures might capture distinct facets of emotion regulation. More broadly, our results highlight concrete methodological challenges for fMRI research on individual differences, with implications extending beyond emotion regulation to the neuroscience of personality, psychopathology, and general well-being.

## Introduction

Emotion regulation is a core determinant of mental and physical health (Sheppes et al., 2015) (Sheppes et al., 2015). It encompasses the strategies people use—intentionally or automatically—to influence their emotional experience (Braunstein et al., 2017; Thompson, 1994). Among these, cognitive reappraisal—changing the interpretation of a situation to alter its emotional impact—has received particular attention in psychology, psychiatry, and the broader public. It is central to cognitive-behavioral therapy and linked to resilience, lower symptom burden, and improved daily affect (D’Agostino et al., 2017).

Neuroimaging studies have extensively mapped the neural basis of reappraisal. Functional Magnetic Resonance Imaging (fMRI) consistently implicates a fronto-parietal network, including lateral and medial prefrontal regions, often along with reduced amygdala activation during reappraisal of negative stimuli (Buhle et al., 2014; Min et al., 2022; Morawetz et al., 2017; Powers & LaBar, 2019). These findings are based primarily on *within-person* designs that contrast experimental conditions like “reappraise” versus “permit emotion” in the same individuals. Yet, many studies — and broader theories — extend these neural within-person mechanisms to explain *between-person* individual differences in well-being or psychopathology (Picó-Pérez et al., 2017; Sicorello & Schmahl, 2021), which is not necessarily valid.

Inferences from within- to between-person levels often fall prey to the ecological fallacy (Kievit et al., 2013). This common phenomenon occurs when within- and between-person associations do not have the same causal structure (Rohrer & Murayama, 2023). For instance, people who habitually use reappraisal tend to report better mental health than other people (between-person level; comparison between individuals). Yet in everyday life, reappraisal often occurs in response to situational stressors and may therefore coincide with a worse mood in that moment (within-person level; comparison between conditions). Similarly, psychologically meaningful individual differences in neurovascular factors — such as fitness or age — can influence the strength of global fMRI responses, potentially leading to ecological fallacies and complicating comparisons between people (Fabiani et al., 2014; Sicorello et al., 2025).

Compounding this issue, regions activated during reappraisal also subserve other functions, making it unclear whether they reflect regulation capabilities per se (Kragel et al., 2018; Poldrack, 2011).In most studies, their relation to self-reported emotion regulation capabilities is not tested directly, even on a within-person level (though see: Bo et al., 2024; Ochsner et al., 2002; Wager et al., 2008). Moreover, task designs may not mirror naturalistic real-life regulation, raising questions about their validity and practical relevance for important individual differences (Enkavi & Poldrack, 2020; McDermott et al., 2018; Powers & LaBar, 2019; Sicorello et al., 2025). Meta-analyses on reappraisal in mental disorders have yielded mixed results, with inconsistent findings for commonly implicated regulatory regions like the dorsolateral prefrontal cortex (dlPFC; Khodadadifar et al., 2022; Morawetz et al., 2025; Sicorello & Schmahl, 2021). A recent systematic review found partial convergence in the hypothesized fronto-limbic circuits, but studies were underpowered (median N = 25), heterogeneous, and too few for formal meta-analysis (Morawetz & Basten, 2024).

To address the uncertainty surrounding the neural correlates of individual differences in reappraisal capabilities, we conducted a pre-registered, well-powered analysis of 40 task-based fMRI datasets (N = 2,175; Table S1), aggregated by the newly founded Neurobiology of Individual Differences in Emotion Regulation (NIDER) consortium. We examined three widely used indices of reappraisal capability (Dörfel et al., 2020): (1) self-reported habitual reappraisal (*trait questionnaires*), (2) task-based reductions in negative affect (*task-based affective ratings*), and (3) task-based reductions in amygdala activation (*amygdala down-regulation*). This enabled us to ask whether these indices reflect a shared underlying core construct—and whether individual differences in reappraisal capability, as indexed by each measure, are systematically associated with neural activity in a priori defined regulatory neural networks.

## Results

### Convergence between reappraisal capabilities from trait questionnaires, task-based affective ratings and amygdala down-regulation

In a first step, we aimed to assess the overlap between the three main outcome measures of cognitive reappraisal capabilities. If these three measures represent a similar core construct they should show a meaningful association. There was a small statistically significant association between amygdala down-regulation and task-based affective ratings (Table 1). As hypothesized, participants with stronger down-regulation of the amygdala via cognitive reappraisal also showed a better down-regulation of task-based affective ratings. Yet, the effect size was comparatively small. To detect this effect size with a statistical power of 80%, a single study would need to test 1105 participants. There was no significant association between trait questionnaires and either outcome, i.e., amygdala down-regulation or task-based affective ratings. The overlap (shared variance) between the three outcome measures is shown in Figure 1, which indicates no meaningful overlap between all three outcomes.

**Table 1.**
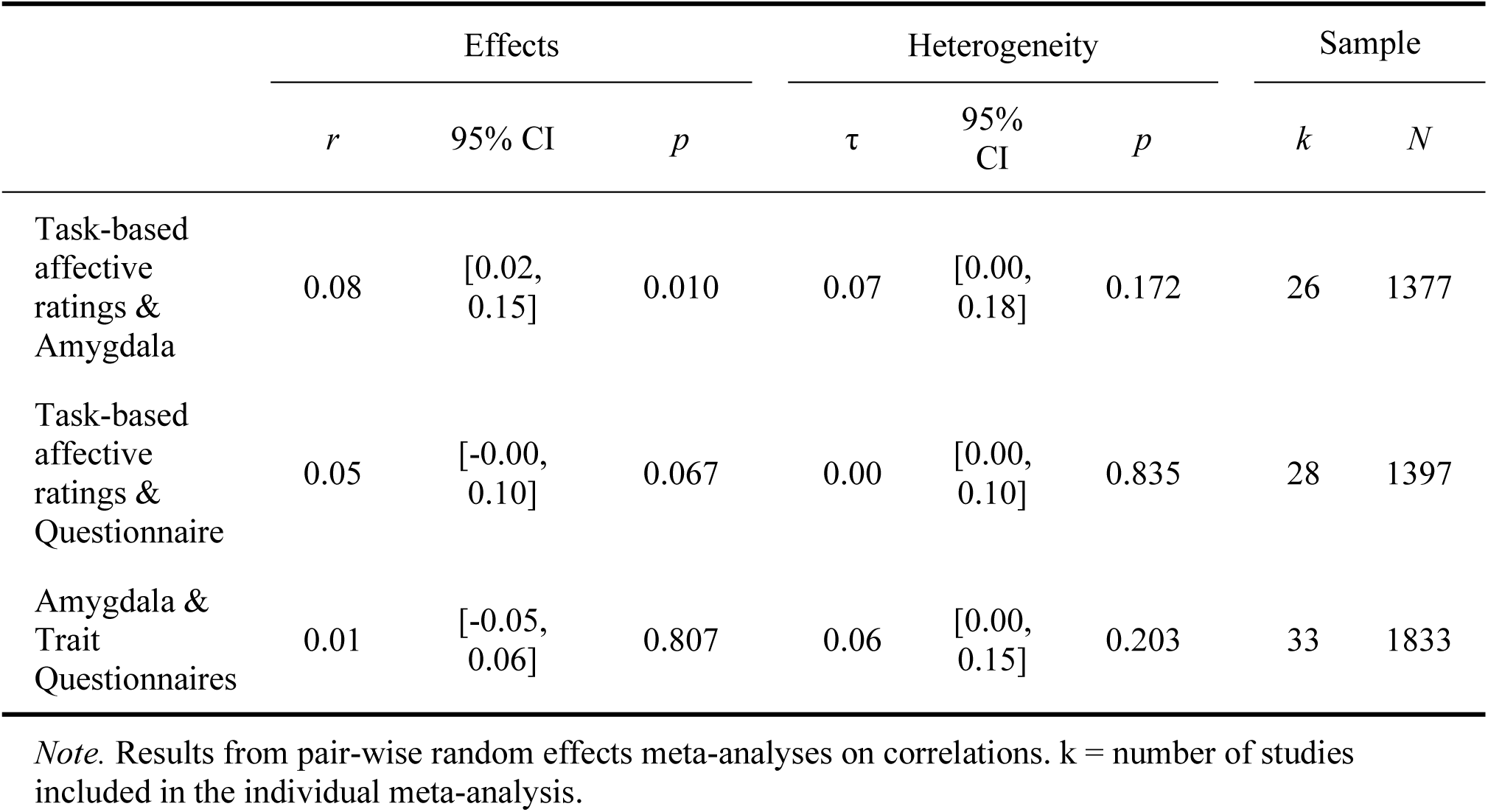
Meta-analytic associations between task-based affective ratings, trait questionnaires, and amygdala down-regulation.

**Figure 1.**
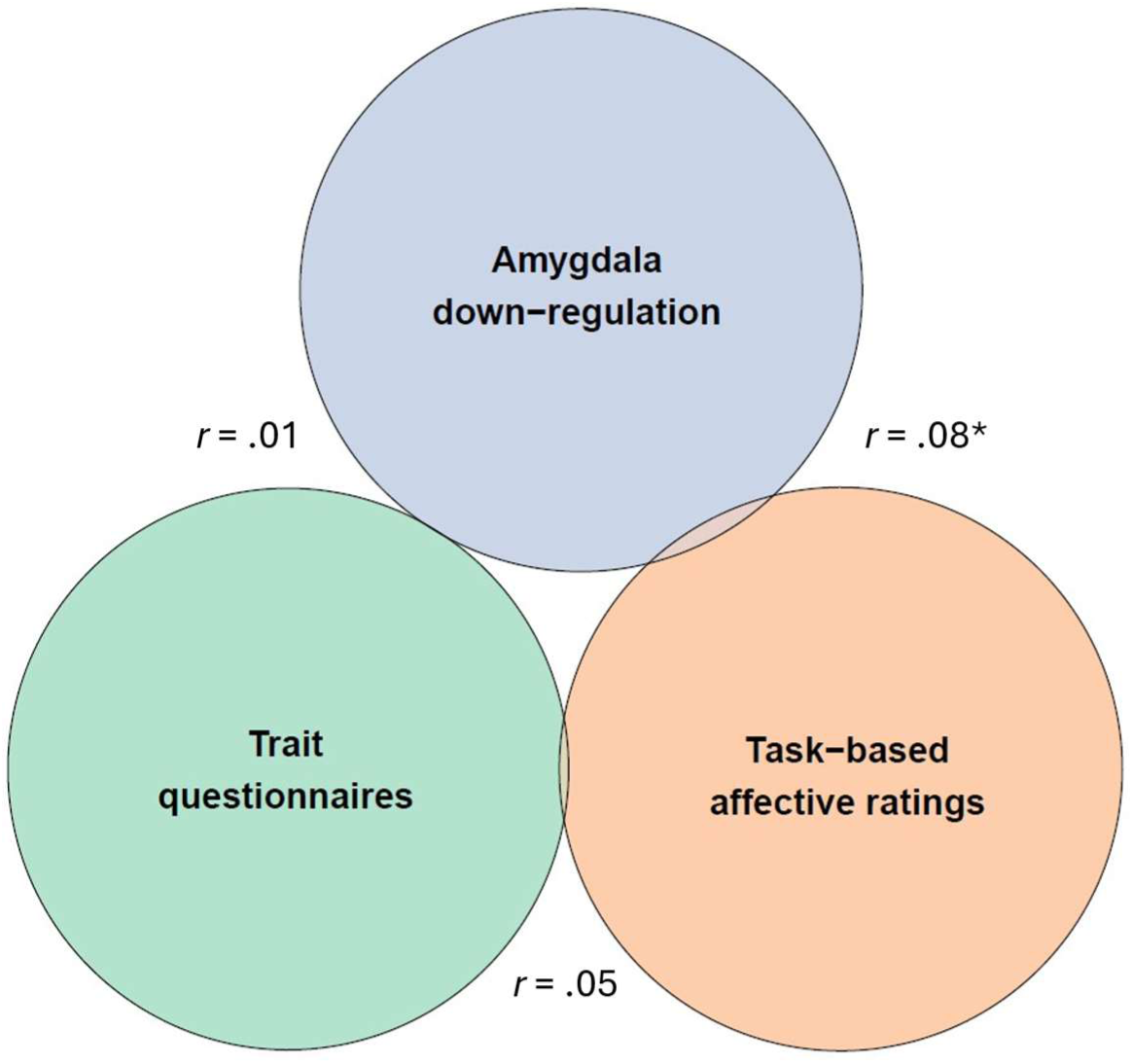
Overlap (shared variance) between the three outcome measures together with the corresponding meta-analytic correlations between pairs of outcomes.

Between-study heterogeneity was not statistically significant for all three tests and was particularly small for the non-significant correlations, indicating these null findings are unlikely to be explained by differences in study design. Forest plots are shown in Figures S1-S3.

### Associations with regulatory neural networks

#### Trait questionnaires

There was no statistically significant association between trait questionnaires and neural activity during reappraisal. The correlations with average activity in the two predefined networks were negligibly small (both |*r*| ≤ .04, *p* > .18, uncorrected; *k* = 37, *N* = 2091) and no voxels survived the correction for multiple comparisons in the voxel-wise network of interest or the whole-brain approach. All brain-wide effect sizes were smaller than |r| ≤ .11. These null results were stable when using a jackknife procedure, i.e., repeating the meta-analysis while leaving out one study at a time. For separate analyses on single studies, only three had significant voxels after FDR correction, with relatively low numbers, which is still expected by chance with the current number of studies (Gianaros et al. [2020]: 2 voxels; Dörfel et al. [2014]: 9 voxels; Sandner et al. [2021]: 22 voxels).

#### Task-based affective ratings

As for trait questionnaires, there was no statistically significant association between average down-regulation of task-based affective ratings and average neural activity during reappraisal in the preregistered regulatory networks (both |r| < .03, p > .22; *k* = 31, *N* = 1958). No voxels within these networks survived correction for multiple comparisons in voxel-wise analyses. Nevertheless, outside of the regulatory network there was a significant negative association in a cluster of 86 voxels. The study by Brehl and colleagues (2021) strongly influenced the results according to the jackknife procedure (Figure S4). Its exclusion led to a substantial increase to 586 significant voxels, mainly covering regions in the somatomotor and dorsal attention network, but also smaller clusters in the visual cortex, caudate nucleus and brainstem (Figure 2C, Table S2). The association between these brain regions and task-based affective ratings was negative, indicating better down-regulation of negative emotions was associated with decreased activations in these areas (average *r* = -.11, range = -.16–-.09). Neurosynth decoding suggested the meta-analytic activation map is most similar to language processing and most dissimilar to motoric activations.

**Figure 2.**
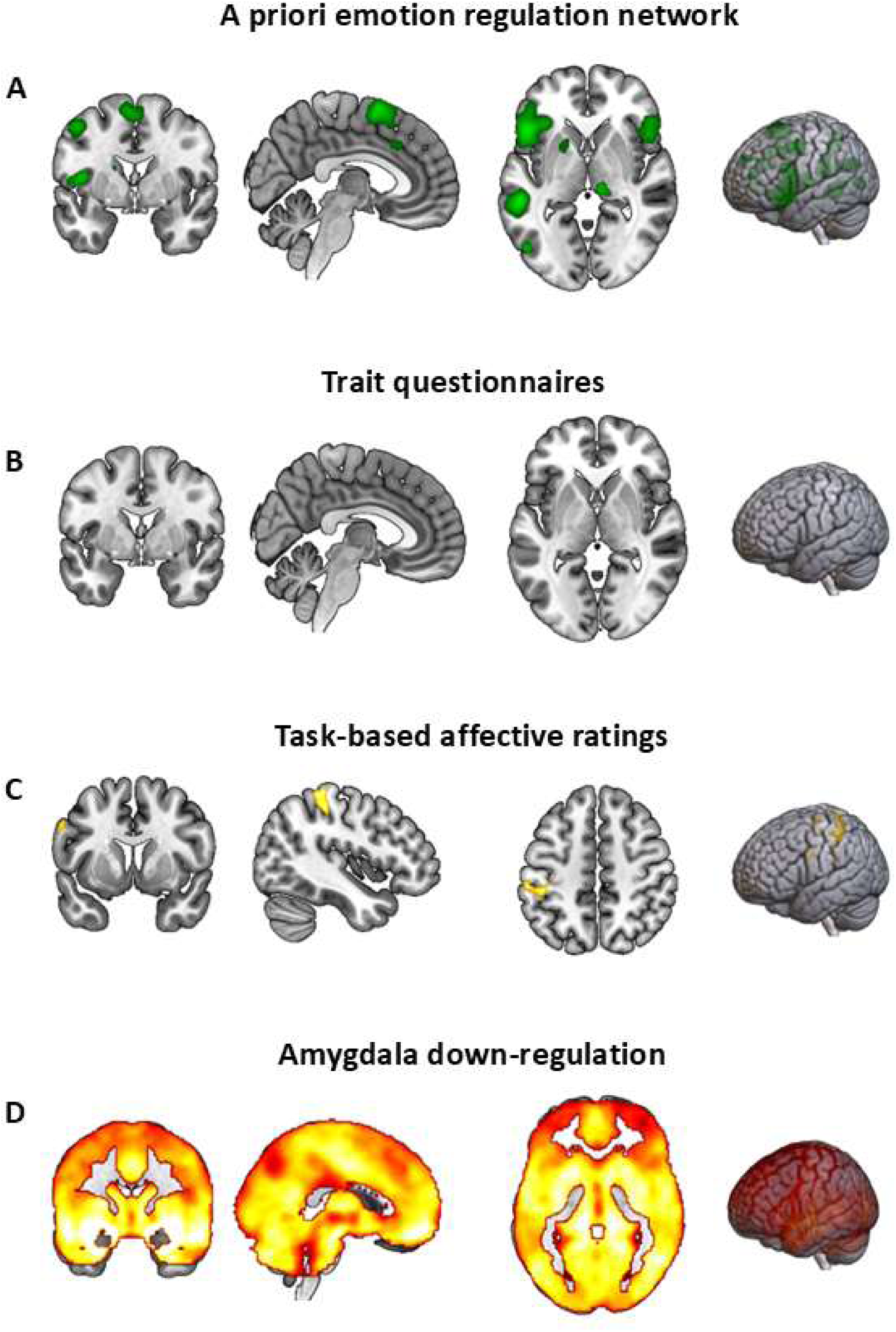
(A) A priori emotion regulation networks of interest based on a previous meta-analysis (Morawetz et al., 2020). (B-D) Significant brain-wide associations between activation in the contrast [reappraise-view] and the three main outcomes for reappraisal capabilities. All significant correlations had negative signs. Therefore, brighter colours correspond to larger *negative* associations. MNI coordinates for slices were: (A) x = 4, y = 0, z = 0; (B) x = 4, y = 0, z = 0; (C) x = −44, y = 7, z = 4; (D) x = 0, y = 0, z = 0.

We repeated the whole-brain analyses separately for each single study. There were 151 significant voxels in Brehl et al. (2021) and 276 significant voxels in Gianaros et al. (2020). While it is still within chance-based expectations to have two out of 28 studies show significant results, it is still noteworthy that these were the two studies with the largest sample sizes (242 and 176 respectively). Moreover, these results included many regions which are part of networks implicated in cognitive reappraisal and emotion processing, such as the amygdala, the hypothalamus, the ventral and dorsal attention networks as well as a fronto-parietal network, but also the somatosensory cortex and the default mode network (Bo et al., 2024).

#### Amygdala down-regulation

A person’s ability to either *down*-regulate their amygdala or *up*-regulate regulatory regions is often interpreted as an indicator of interindividual reappraisal capability. Such indices have been criticized as there is evidence that people differ in their global brain responses, likely due to biological confounders (Fabiani et al., 2014), which might dominate any specific regional effects and dilute their usefulness as a marker of an individual’s emotion regulation capability (Sicorello et al., 2025). In the case of reappraisal, this might be particularly problematic as opposite effects are expected for regulatory regions and emotion generating regions like the amygdala.

In support of this criticism, across studies, participants with stronger amygdala responses during reappraisal also had substantially stronger responses in the rest of the brain in comparison to other participants (Figure S5; mean *r* = .27, range =-.08–.68). This included the two regulatory networks (first network: *r* < .29, *p* < .001; second network: *r* < .24, *p* < .001; *k* = 35, *N* = 2088) and generally most parts of the brain, with significant positive correlations for over 170,000 voxels versus only one voxel with a negative correlation (Figure 2D). Hence, a person is unlikely to simultaneously show both lower amygdala responses and higher regulatory network responses (e.g., in the dlPFC) when compared to other people using fMRI. This limits the interpretational validity of single specific brain regions for interindividual differences in reappraisal capabilities, especially when hypothesizing effects in different directions, as is the case for the amygdala and regulatory regions.

In an attempt to test the specificity of any relevant target regions, we tested whether this finding generalizes to other brain regions beyond the amygdala by calculating the average pairwise between-person correlation for neural responses during reappraisal in 470 anatomical regions (Wager, 2024) of a publicly available dataset (*N* = 33; Wager et al., 2008). This showed that responses of any two regions across the brain are generally highly correlated on a between-person level (*r* = .58, *SD* = .18), limiting their spatial specificity and the inferential value of this approach. Hence, if a person has a stronger fMRI response in one brain region than other people, that person is likely to have a stronger fMRI response in any other region as well. This represents a phenomenon that has been previously noted (Jabakhanji et al., 2022), yet commonly overlooked in the study of individual differences, including most research in the domain of well-being and psychopathology.

We further tested whether these high between-person correlations of different brain regions are specific to cognitive reappraisal or generalize across other psychological domains. We again calculated the average pairwise between-person correlation for activity in the 470 anatomical regions in an openly available multi-task dataset (Kragel et al., 2018). This dataset combines 18 studies across the domains of negative emotion, pain, and cognitive control, each represented by three subdomains, in turn each represented by two studies (*n* = 15 per study, total *N* = 270]). Overall, we found the same pattern of overly large bivariate between-person correlations across the whole brain (mean correlation across domains: *r* = .37; Figure 3).

**Figure 3.**
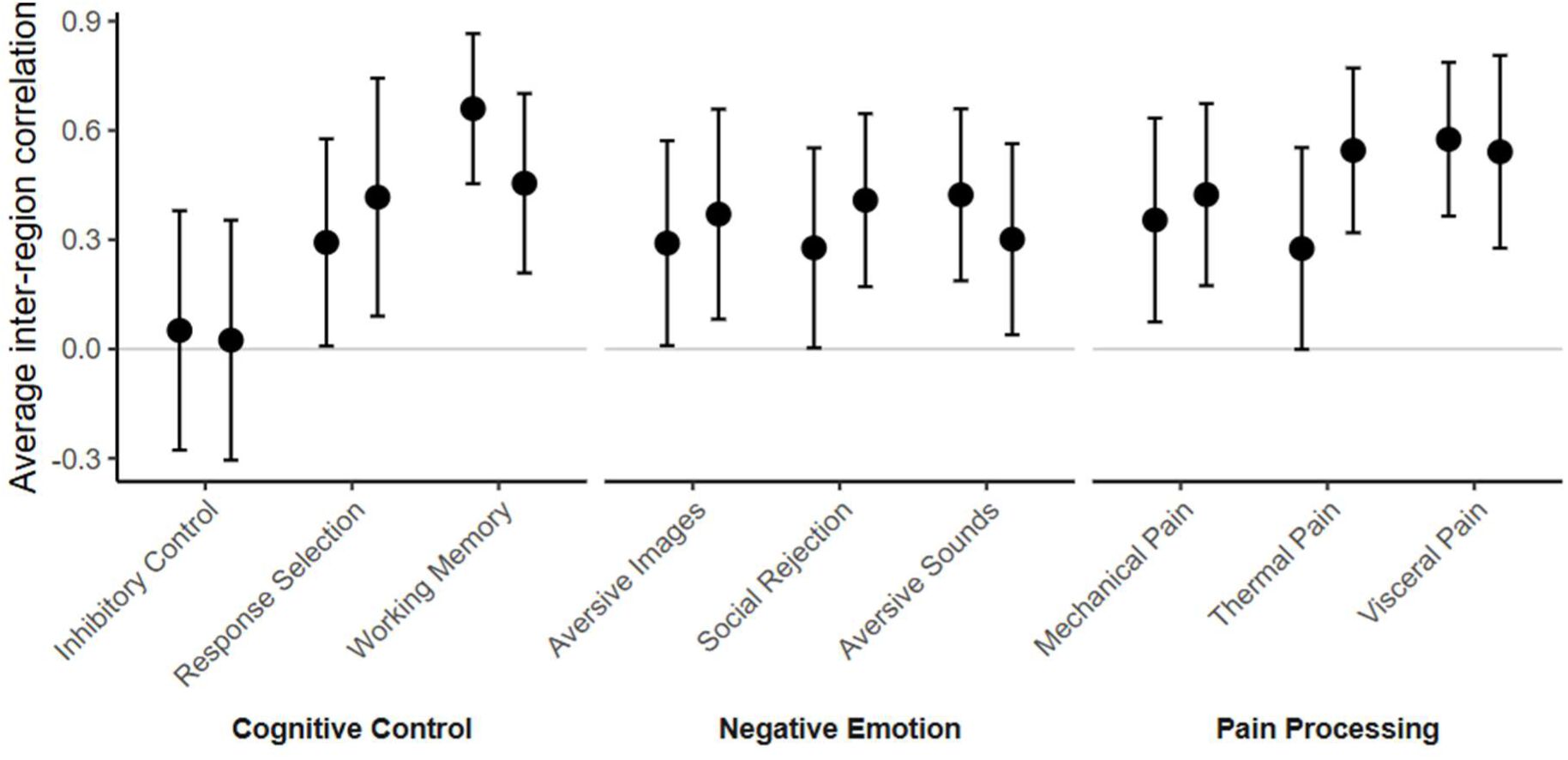
Between-person correlations for pairs of regions based on an open dataset with three domains, three subdomains per domain, and two studies per subdomain (*n* = 15 per study, total *N* = 270). Each estimate represents the average correlation between all possible pairs of 470 regions. Whiskers represent the standard deviation.

In sum, this suggests that interindividual differences in specific regional brain responses, like the amygdala or the dlPFC, are strongly confounded by interindividual differences in global signal responses with implications beyond emotion regulation.

### Statistical power

We computed statistical power for a range of correlation effect sizes as a function ofbetween-study heterogeneity (*τ*), the number of voxels corrected for with the FDR procedure (two networks, four networks, and whole-brain), and the number of voxels assumed to be true positives (corrected for spatial dependence, for methodological details and validation of the procedure see supplements). Sample sizes were fixed to the values used for the whole-brain correlations with questionnaires. Results are shown in Figure 4. Given a conservative minimum number of 100 true positive voxels, we had sufficient statistical power to detect small correlations between 0.05-0.2, depending on the heterogeneity between studies. Heterogeneity was overall very low in all conducted analyses, indicating the statistical power is likely sufficient for the lower end of these effect sizes.

**Figure 4.**
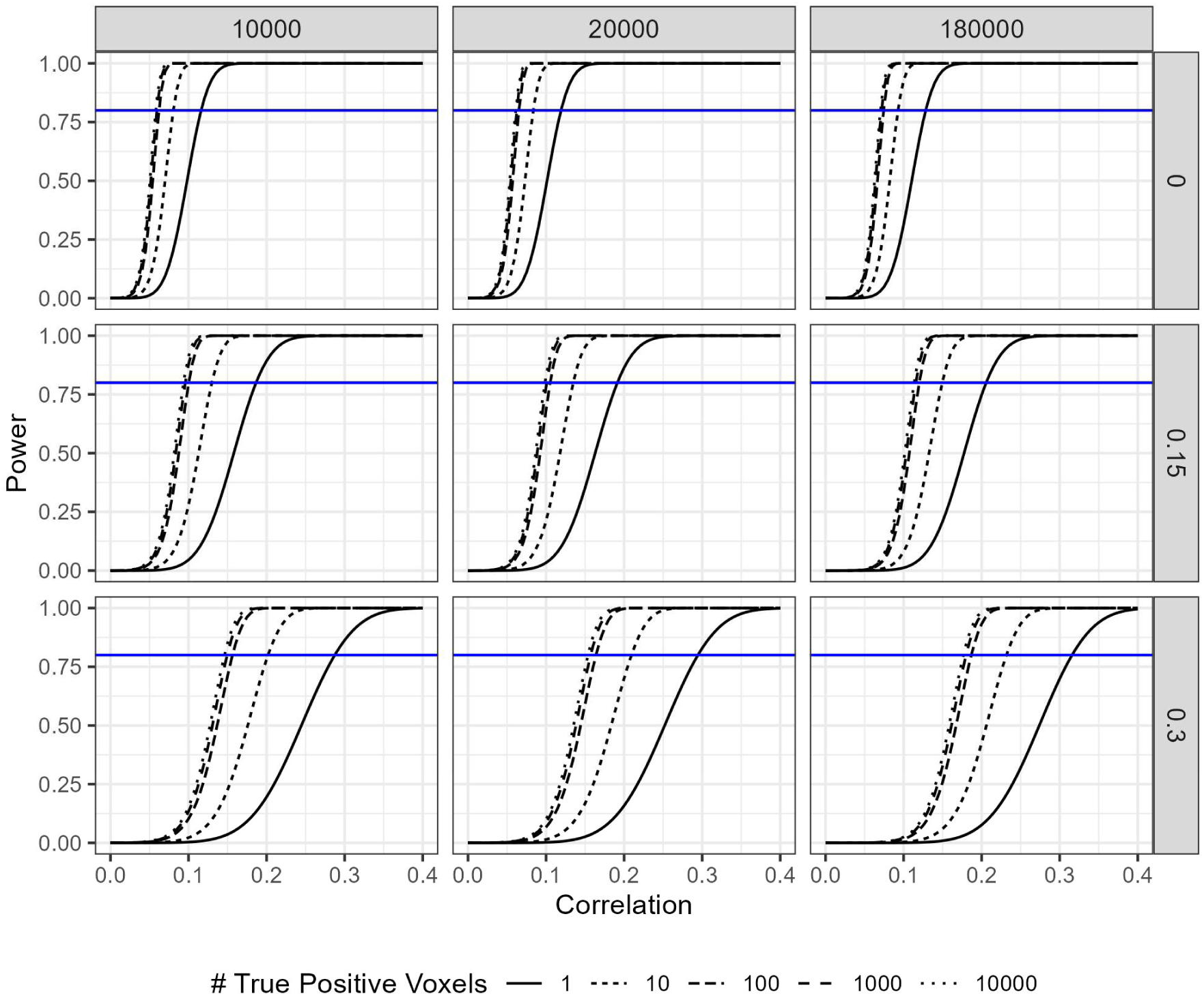
Power analyses for a range of hypothetical parameters. Blue lines indicate 80% power. Rows show results for different values of between-study heterogeneity (*τ*); columns show the number of tests corrected based on the two preregistered networks of interest, an expanded network of interest using the full results of Morawetz et al. (2020), and whole-brain data after masking for gray matter and a minimum number of 20 studies with valid data. The legend shows the number of voxels assumed to be true positives. Each of these numbers corresponds to a smaller (analytically derived) number of effective (i.e., independent) tests, which were used as input in the power analysis (1=1, 10=7, 100=34, 1000=45, 10000=52).

## Discussion

Despite extensive research on the neural basis of cognitive reappraisal, the association between these neural measures and individual differences in psychologically assessed reappraisal capability remain inconsistent. Studies on well-being and psychopathology often interpret reduced prefrontal activation (e.g., in the dlPFC) as reflecting impairments in cognitive reappraisal, yet recent meta-analyses show little convergence of such group differences across or within disorders in frontal regulatory regions (Khodadadifar et al., 2022; Morawetz et al., 2025; Sicorello & Schmahl, 2021). Dimensional approaches—linking brain function to measures of reappraisal capabilities—remain rare and typically underpowered. To overcome this, we conducted a large-scale, preregistered consortium analysis with a sample approximately 20 times larger than the largest sample reported in a recent systematic review (Morawetz & Basten, 2024). By pooling whole-brain data from diverse studies, most of which had not previously performed the targeted analyses, we aimed to identify robust neural markers of reappraisal capabilities and contribute to a better understanding of neurobiological individual differences.

There was a striking lack of convergence between the three indices commonly used to indicate reappraisal capabilities: trait questionnaires, task-based affective ratings, and amygdala down-regulation. Only the association between the latter two—simultaneously measured task-based affective ratings and amygdala down-regulation—was statistically significant. The small effect size indicates thatmore than 1000 participants are needed for sufficient power, which ismore than 10 times larger than those usually seen in the current literature (Morawetz & Basten, 2024). The low between-study heterogeneity indicates that these results are not due to differences in the included study designs. These findings suggest that the three common indices of reappraisal capability capture qualitatively different facets of reappraisal, despite often being interpreted interchangeably. The absence of a common core among these indicators challenges the assumption that they reflect a shared underlying capacity.

Similarly, there were no brain-behaviour associations between either trait questionnaires or task-based affective ratings and neural activity in established emotion regulation networks. However, for task-based affective ratings, we observed significant associations with lower activity in left sensorimotor and dorsal attention networks—regions not commonly found in reappraisal research. The lower responses in dorsal attention networks might relate to a lowered focus on negative external stimuli (Vossel et al., 2014), while the lowered sensorimotor responses might relate to the embodied aspects of emotional experiences (Reddan et al., 2024). Neurosynth decoding also indicated an increased language-related activation, consistent with the verbal and cognitive nature of reappraisal strategies. Together, these results suggest that regions outside canonical process-based emotion regulation networks—including the prefrontal cortex—may better reflect individual differences in regulation capability, albeit with still very small effect sizes.

The disconnect between trait questionnaires and task-based affective ratings offers a partial plausible explanation for the null finding for the association between trait questionnaires and task related brain activity. If questionnaires are not matched well to tasks on a behavioral level, meaningful associations between trait questionnaires and neural activity during these tasks seem less likely. The studies aggregated here predominantly used the most common reappraisal task (Ochsner et al., 2002) and the two most common reappraisal questionnaires (Emotion Regulation Questionnaire [ERQ; Gross & John, 2003]; and Cognitive Emotion Regulation Questionnaire [CERQ; Jermann et al., 2006]). Therefore, these results have implications for a large body of research. Both questionnaires measure habitual tendencies, rather than ability. This habitual tendency or frequency of using reappraisal is only moderately correlated with reappraisal abilities in naturalistic daily life settings (Koval et al., 2023). Correlations of these trait questionnaires with measures from experimental settings are even lower (Ford et al., 2017), as we have shown here. Regarding tasks, there has been criticism that, for example, distancing oneself from pictorial stimuli by imagining it as not real or being viewed from a larger distance (common strategy instructions), might lack ecological validity for reappraisal outside the laboratory (Powers & LaBar, 2019). Hence, more attention should be given to the exact aspect of emotion regulation operationalized in a task or a questionnaire, with a pressing need for novel task designs and more naturalistic stimuli. This is not restricted to emotion regulation, but has also been recently shown for emotion *generation* and a large range of related trait questionnaires measuring negative affectivity (Sicorello et al., 2025).

Importantly, a mismatch between tasks and questionnaires does not explain the very small associations between task-based ratings and neural measures. In principle, within-person associations between brain responses and task-based self-reports can be very robust (Chang et al., 2015; Zhou et al., 2021). In contrast, between-person associations for person-wise averages of the same self-reports appear to have a ceiling around r = .20-.30, even with machine learning approaches (Gianaros et al., 2022; Han et al., 2022; Sicorello et al., 2025). One reason may be that standard tasks were designed for process-based research and do not sufficiently elicit relevant individual differences, for example, because they are not sufficiently naturalistic (Hedge et al., 2018). Another reason is that people appear to have very different average whole-brain responses to tasks, related to biological confounds like neurovascularity or noise sources such as MR-system related fluctuations and subject movement (Fabiani et al., 2014; Huber et al., 2024; Liu, 2016; Sicorello et al., 2025; Ward et al., 2020). We found that fMRI responses during reappraisal were highly correlated across regions on a between-person level, including the amygdala and prefrontal cortex. This pattern extended beyond reappraisal to multiple task domains, suggesting that person-level brain-wide responsivity severely limits the interpretational validity of specific regional effects. Accounting for this dependency in the fMRI signal for such brain-wide responses is likely a crucial analytic step towards larger and more valid effects (Davis et al., 1998; Sicorello et al., 2025). Yet, as the lively debate around global signal regression in resting state fMRI research has made clear, such approaches are anything but universally accepted (Murphy & Fox, 2017).

Our study focused on non-clinical samples, as these most often consist of case-control designs with complex groups concerning their symptom and medication profiles. A representative sample would be expected to have 1.3-1.5 times the standard deviation in the ERQ compared to those studies we included (Preece et al., 2020). While a representative sample with this range would lead to only slightly increased effect sizes, the focus on transdiagnostic samples that cover large parts of clinical dimensions could provide important extensions of our study. Moreover, most studies in the literature use the same questionnaires, which was reflected in our study. Questionnaires focusing on reappraisal *ability*, rather than habitual use, might lead to slightly larger associations.

We provide a reusable implementation of image-based meta-analyses together with accessible power analyses tools, which might aid similar projects to consolidate the large body of fMRI research beyond emotion regulation. Most fMRI studies employ a large range of questionnaires for individual differences, which are usually not tested for brain-wide associations and therefore represent an extremely valuable data resource. Using a meta-analytic approach has similar statistical properties as random effects mega-analysis on individual-level data (Eisenhauer, 2021; Koile & Cristia, 2021) and provides an efficient tool to retain and quantify the heterogeneity between studies, leading to more generalizable results. Nonetheless, statistical tools alone are insufficient. Our findings challenge widespread assumptions about the neural correlates of individual differences in cognitive reappraisal and call for a more conscious and explicit alignment between constructs, measurements, and analytic levels.

In summary, our findings challenge prevailing assumptions about the neural correlates of interindividual capabilities in cognitive reappraisal and raise fundamental questions about how emotion regulation is measured across levels of analysis. The weak convergence between common outcome measures and their inconsistent associations with canonical reappraisal networks underscore the need for better-aligned behavioral, trait-based, and neural constructs for them to capture a shared core of cognitive reappraisal. This is especially relevant given that extremely large-scale resting-state efforts have failed to reveal replicable neural correlates of emotion-related traits (Marek et al., 2022; Schulz et al., 2024). Employing tasks that are well-designed to elicit the individual differences of interest is a crucial strategy for a better neurobiological understanding of emotional well-being (Sicorello et al., 2025). Still, we demonstrated that interindividual differences in whole-brain responses during tasks are a methodological issue that likely affects the vast majority of the task-based fMRI literature, beyond emotion regulation. Ultimately, identifying robust and generalizable neural markers of self-regulation requires both methodological progress and conceptual clarity in defining what we seek to measure (Brandt & Mueller, 2022; Sicorello et al., 2025). Until then, we should refrain from strong statements concerning a person’s emotion regulation capabilities based on their prefrontal cortex, amygdala, or any other specific brain region or network.

## Methods

### Consortium procedures

Studies were eligible for inclusion if they measured (1) task-based fMRI (2) during a cognitive reappraisal task (3) using negative stimuli (4) to compare a down-regulation condition with a maintaining/permitting condition (5) and included a trait-like questionnaire measure for cognitive reappraisal (6) in a healthy sample. The latter criterion was used to avoid excessive heterogeneity in disorder presentation and medications (see the discussion section). Contributing research groups were invited using two strategies. First, we conducted a systematic literature search, contacting 64 authors via e-mail or Researchgate. Second, we used mailing lists of academic societies to advertise the consortium. In total, this led to 32 contributing research teams with 40 datasets comprising up to 2,175 participants. For an overview of study characteristics see Table S1. Nine studies were unpublished and none of the studies performed the relevant analyses for previous publications. Most papers used the ERQ to measure reappraisal, while six used the CERQ and only one study used a different questionnaire (FEEL-E). The majority of studies used pictorial stimuli, two studies used videos, one study used autobiographical memory cues, and one study used Cyberball.

### Literature search

We conducted a systematic literature search of articles using the PubMed, Web of Science, and PsychINFO databases. We searched for the following keywords: “*emotion regulation”* cross referenced with “*fMRI”* or “*neuroimaging”* or “*functional magnetic resonance imaging”* or “*functional MRI”*. This search process yielded a total of 2,633 potentially relevant articles on July 7, 2021 (after duplicates were removed). Two independent reviewers (JZ, MS) systematically examined titles and relevant abstracts using the Rayyan website (Ouzzani et al., 2016) to determine whether an article fulfills the criteria to be screened. The following criteria were applied: The study included original empirical results, was written in English or German, included adult healthy participants, and assessed an explicit emotion regulation paradigm during fMRI measurement. In a next step, for all remaining studies full texts were screened by three independent reviewers (JZ, MB, DL) and they were included if (1) participants were told to use reappraisal strategies to modulate a negative emotion, (2) an emotion regulation questionnaire was assessed (e.g., ERQ), (3) a control condition was included in which participants were confronted with emotional stimuli but did not regulate their emotions, and (4) negative stimuli were used in the emotion regulation task. Finally, a total of *k* = 61 studies fulfilled all inclusion criteria. Relevant articles in the authors’ library were also reviewed for titles that might have been missed by our literature search. Studies identified in this manner (*k* = 3) were collected for inclusion. Of the *k* = 64 studies, *k* = 17 reported the correlation between fMRI BOLD activation (‘reappraise-view’) and emotion regulation questionnaire data. The authors of all 64 studies were contacted. See Figure S6 for a flow chart depiction of the screening and selection of studies via literature search.

### Data extraction

Research groups prepared up to three unthresholded group-level statistical images, containing voxel-wise *t*-statistics for the between-person correlation of the whole-brain neural response in the [reappraise - view] contrast and the three outcomes of interest. These outcomes were (1) trait questionnaires of cognitive reappraisal, (2) the person-wise averaged decrease of task-based affective ratings during reappraisal versus view conditions, and (3) and person-wise averaged decrease of amygdala responses during reappraisal versus view conditions.

A homepage with detailed instructions and a Matlab example analysis script was provided to support analyses. Contributors were instructed to compute the three measures of reappraisal capabilities so that higher values correspond to better reappraisal capabilities. For task-based affective ratings, this entailed (re-)coding for higher values to correspond to a more negative emotional response and calculating person-wise averaged down-regulation of affect from the contrast [view - reappraise]; hence, a greater reduction in negative affect via reappraisal leads to higher values. Similarly, for amygdala down-regulation, average activity was extracted from the [view - reappraise] contrast, as higher values correspond to a better down-regulation of the amygdala for sign congruence with the other two outcomes. A standardized amygdala-mask was provided for all research groups based on the intersection of an anatomical amygdala mask (SPM anatomy toolbox v2.2b) and a functional emotion generating network from an emotion regulation meta-analysis (Morawetz et al., 2020), which ensures the anatomical and functional specificity of the amygdala region of interest.

Furthermore, contributors provided direct bivariate correlations between the three reappraisal outcomes (e.g., trait questionnaires and task-based affective ratings) as well as demographic and other study information (Table S1).

Images were quality controlled individually by plotting t-values on a brain template, histograms, and performing brain-wide thresholding at FDR = .05 to check for plausibility. Collectively, quality was controlled by checking for outlier studies within the same emotion regulation outcome category (e.g., mean whole-brain activation, Mahalanobis distance, and covariance matrices). Two images were removed from whole-brain analyses due to insufficient coverage (Berboth et al., 2021; Morawetz et al., 2016). Effective sample sizes for tests are reported in the results section.

### Statistical Analyses

First, we assessed the concordance between the three outcome measures (trait questionnaires, task-based ratings, amygdala down-regulation) using random effects meta-analysis implemented in the R package *meta* using the DerSimonian and Laird approach to estimate between-study heterogeneity (Balduzzi et al., 2019).

Second, we tested associations between the three outcomes and two neural emotion regulation networks based on a recent process-based meta-analysis on emotion regulation (Morawetz et al., 2020; networks 1 and 2; Figure 2). These networks are thought to be predominantly implicated in the cognitive control (versus emotion generating) aspects of emotion regulation and mostly include regions in the prefrontal, parietal, and temporal cortex. Images were resampled to the same spatial resolution of 2×2×2 mm³. The statistical core procedure was done in three pre-registered stages. In the first stage, we tested the correlation between average activity in the two networks of interest, respectively, and reappraisal capabilities, with Bonferroni corrections for two tests. In the second stage, we performed voxel-wise tests in the combined two networks with FDR correction based on the Benjamini-Hochberg procedure (FDR = 0.05, ≈10,000 voxels). In a third stage, we repeated the second-stage analysis on the whole brain (≈180,000 voxels). Between-study heterogeneity was tested for statistical significance based on whole-brain data and corrected with the same FDR procedure.

To perform the image-based fMRI meta-analysis, we developed custom Matlab code which interfaces with the fMRI-specific functionality of the CanlabCore toolbox (Wager, 2024). Our procedure converts study-wise unthresholded t-maps to correlation maps and therefore, together with sample size information, offers voxel-wise estimates of effect size, statistical significance, and between-study heterogeneity via the DerSimonian and Laird approach (Borenstein et al., 2009; Bossier et al., 2019). The code was validated by testing congruence with meta-analytic packages in R as well as performing a small meta-analysis on six openly available group-level images of physical pain, which detected typically involved brain regions and was successfully decoded as “pain” using neurosynth (see supplement “Testing the custom meta-analytic functions”, Figure S7 for details).

### Open Science Practices

Aims, literature search strategy, and statistical analyses were pre-registered on PROSPERO: https://www.crd.york.ac.uk/PROSPERO/view/CRD42021243155. Group-level maps and code are openly provided on https://github.com/MaurizioSicorello/NIDER_project. We provide reusable functions to conduct image-based fMRI meta-analyses in Matlab (v2023b).

A notable error in the preregistration was the statement that only studies using pictures as stimuli are included. Rather, the meta-analysis included all types of stimuli, which we stated correctly on the accompanying homepage and the mailing list advertisements. Regardless of this error, the overwhelming majority of studies used pictures as stimuli (88%) and the low statistically non-significant between-study heterogeneities as well as the jackknife procedures indicate that stimulus type does not influence the results.

## References

Balduzzi, S., Rücker, G., & Schwarzer, G. (2019). How to perform a meta-analysis with R: A practical tutorial. Evidence-Based Mental Health, 22(4), 153–160. 10.1136/ebmental-2019-300117

Bo, K., Kraynak, T. E., Kwon, M., Sun, M., Gianaros, P. J., & Wager, T. D. (2024). A systems identification approach using Bayes factors to deconstruct the brain bases of emotion regulation. Nature Neuroscience, 27(5), 975–987. 10.1038/s41593-024-01605-7

Borenstein, M., Larry V., H., Julian P.T., H., & Hannah R., R. (2009). Introduction to Meta-Analysis. John Wiley & Sons, Ltd.

Bossier, H., Nichols, T. E., & Moerkerke, B. (2019). Standardized Effect Sizes and Image-Based Meta-Analytical Approaches for fMRI Data. bioRxiv, 1–61. 10.1101/865881

Brandt, A., & Mueller, E. M. (2022). Negative affect related traits and the chasm between self-report and neuroscience. Current Opinion in Behavioral Sciences, 43, 216–223. 10.1016/j.cobeha.2021.11.002

Braunstein, L. M., Gross, J. J., & Ochsner, K. N. (2017). Explicit and implicit emotion regulation: A multi-level framework. Social Cognitive and Affective Neuroscience, 12(10), 1545–1557. 10.1093/scan/nsx096

Buhle, J. T., Silvers, J. A., Wager, T. D., Lopez, R., Onyemekwu, C., Kober, H., Webe, J., & Ochsner, K. N. (2014). Cognitive reappraisal of emotion: A meta-analysis of human neuroimaging studies. Cerebral Cortex, 24(11), 2981–2990. 10.1093/cercor/bht154

Chang, L. J., Gianaros, P. J., Manuck, S. B., Krishnan, A., & Wager, T. D. (2015). A sensitive and specific neural signature for picture-induced negative affect. PLoS Biology, 13(6), 1–28. 10.1371/journal.pbio.1002180

D’Agostino, A., Covanti, S., Rossi Monti, M., & Starcevic, V. (2017). Reconsidering emotion dysregulation. Psychiatric Quarterly, 88(4), 807–825. 10.1007/s11126-017-9499-6

Davis, T. L., Kwong, K. K., Weisskoff, R. M., & Rosen, B. R. (1998). Calibrated functional MRI: Mapping the dynamics of oxidative metabolism. Proceedings of the National Academy of Sciences, 95(4), 1834–1839. 10.1073/pnas.95.4.1834

Dörfel, D., Gärtner, A., & Scheffel, C. (2020). Resting State Cortico-Limbic Functional Connectivity and Dispositional Use of Emotion Regulation Strategies: A Replication and Extension Study. Frontiers in Behavioral Neuroscience, 14. 10.3389/fnbeh.2020.00128

Eisenhauer, J. G. (2021). Meta-analysis and mega-analysis: A simple introduction. Teaching Statistics, 43(1), 21–27. 10.1111/test.12242

Enkavi, A. Z., & Poldrack, R. A. (2020). Implications of the lacking relationship between cognitive task and self report measures for psychiatry. Biological Psychiatry: Cognitive Neuroscience and Neuroimaging. 10.1016/j.bpsc.2020.06.010

Fabiani, M., Gordon, B. A., Maclin, E. L., Pearson, M. A., Brumback-Peltz, C. R., Low, K. A., McAuley, E., Sutton, B. P., Kramer, A. F., & Gratton, G. (2014). Neurovascular coupling in normal aging: A combined optical, ERP and fMRI study. NeuroImage, 85, 592–607. 10.1016/j.neuroimage.2013.04.113

Ford, B. Q., Karnilowicz, H. R., & Mauss, I. B. (2017). Understanding reappraisal as a multi-component process: The psychological health benefits of attempting to use reappraisal depend on reappraisal success. Emotion (Washington, D.C.), 17(6), 905–911. 10.1037/emo0000310

Gianaros, P. J., Rasero, J., DuPont, C. M., Kraynak, T. E., Gross, J. J., McRae, K., Wright, A. G. C., Verstynen, T. D., & Barinas-Mitchell, E. (2022). Multivariate Brain Activity while Viewing and Reappraising Affective Scenes Does Not Predict the Multiyear Progression of Preclinical Atherosclerosis in Otherwise Healthy Midlife Adults. Affective Science, 3(2), 406–424. 10.1007/s42761-021-00098-y

Gross, J. J., & John, O. P. (2003). Individual differences in two emotion regulation processes: Implications for affect, relationships, and well-being. Journal of Personality and Social Psychology, 85(2), 348–362. 10.1037/0022-3514.85.2.348

Han, X., Ashar, Y. K., Kragel, P., Petre, B., Schelkun, V., Atlas, L. Y., Chang, L. J., Jepma, M., Koban, L., Losin, E. A. R., Roy, M., Woo, C.-W., & Wager, T. D. (2022). Effect sizes and test-retest reliability of the fMRI-based neurologic pain signature. NeuroImage, 247, 118844. 10.1016/j.neuroimage.2021.118844

Hedge, C., Powell, G., & Sumner, P. (2018). The reliability paradox: Why robust cognitive tasks do not produce reliable individual differences. Behavior Research Methods, 50(3), 1166–1186. 10.3758/s13428-017-0935-1

Huber, D., Rabl, L., Orsini, C., Labek, K., & Viviani, R. (2024). The fMRI global signal and its association with the signal from cranial bone. NeuroImage, 297, 120754. 10.1016/j.neuroimage.2024.120754

Jabakhanji, R., Vigotsky, A. D., Bielefeld, J., Huang, L., Baliki, M. N., Iannetti, G., & Apkarian, A. V. (2022). Limits of decoding mental states with fMRI. Cortex, 149, 101–122. 10.1016/j.cortex.2021.12.015

Jermann, F., Van der Linden, M., d’Acremont, M., & Zermatten, A. (2006). Cognitive Emotion Regulation Questionnaire (CERQ). European Journal of Psychological Assessment, 22(2), 126–131. 10.1027/1015-5759.22.2.126

Khodadadifar, T., Soltaninejad, Z., Ebneabbasi, A., Eickhoff, C. R., Sorg, C., Van Eimeren, T., Vogeley, K., Zarei, M., Eickhoff, S. B., & Tahmasian, M. (2022). In search of convergent regional brain abnormality in cognitive emotion regulation: A transdiagnostic neuroimaging meta-analysis. Human Brain Mapping, 43(4), 1309– 1325. 10.1002/hbm.25722

Kievit, R. A., Frankenhuis, W. E., Waldorp, L. J., & Borsboom, D. (2013). Simpson’s paradox in psychological science: A practical guide. Frontiers in Psychology. 10.3389/fpsyg.2013.00513

Koile, E., & Cristia, A. (2021). Toward Cumulative Cognitive Science: A Comparison of Meta-Analysis, Mega-Analysis, and Hybrid Approaches. Open Mind, 5, 154–173. 10.1162/opmi_a_00048

Koval, P., Kalokerinos, E. K., Greenaway, K. H., Medland, H., Kuppens, P., Nezlek, J. B., Hinton, J. D. X., & Gross, J. J. (2023). Emotion regulation in everyday life: Mapping global self-reports to daily processes. Emotion, 23(2), 357–374. 10.1037/emo0001097

Kragel, P. A., Kano, M., Van Oudenhove, L., Ly, H. G., Dupont, P., Rubio, A., Delon-Martin, C., Bonaz, B. L., Manuck, S. B., Gianaros, P. J., Ceko, M., Reynolds Losin, E. A., Woo, C.-W., Nichols, T. E., & Wager, T. D. (2018). Generalizable representations of pain, cognitive control, and negative emotion in medial frontal cortex. Nature Neuroscience, 21(2), 283–289. 10.1038/s41593-017-0051-7

Liu, T. T. (2016). Noise contributions to the fMRI signal: An overview. NeuroImage, 143, 141–151. 10.1016/j.neuroimage.2016.09.008

Marek, S., Tervo-Clemmens, B., Calabro, F. J., Montez, D. F., Kay, B. P., Hatoum, A. S., Donohue, M. R., Foran, W., Miller, R. L., Hendrickson, T. J., Malone, S. M., Kandala, S., Feczko, E., Miranda-Dominguez, O., Graham, A. M., Earl, E. A., Perrone, A. J., Cordova, M., Doyle, O.,… Dosenbach, N. U. F. (2022). Reproducible brain-wide association studies require thousands of individuals. Nature, 603(7902), 654–660. 10.1038/s41586-022-04492-9

McDermott, T. J., Kirlic, N., & Aupperle, R. L. (2018). Roadmap for optimizing the clinical utility of emotional stress paradigms in human neuroimaging research. Neurobiology of Stress, 8(November 2017), 134–146. 10.1016/j.ynstr.2018.05.001

Min, J., Nashiro, K., Yoo, H. J., Cho, C., Nasseri, P., Bachman, S. L., Porat, S., Thayer, J. F., Chang, C., Lee, T.-H., & Mather, M. (2022). Emotion Downregulation Targets Interoceptive Brain Regions While Emotion Upregulation Targets Other Affective Brain Regions. Journal of Neuroscience, 42(14), 2973–2985. 10.1523/JNEUROSCI.1865-21.2022

Morawetz, C., & Basten, U. (2024). Neural underpinnings of individual differences in emotion regulation: A systematic review. Neuroscience and Biobehavioral Reviews, 162, 105727. 10.1016/j.neubiorev.2024.105727

Morawetz, C., Bode, S., Derntl, B., & Heekeren, H. R. (2017). The effect of strategies, goals and stimulus material on the neural mechanisms of emotion regulation: A meta-analysis of fMRI studies. Neuroscience and Biobehavioral Reviews, 72, 111–128. 10.1016/j.neubiorev.2016.11.014

Morawetz, C., Hemetsberger, F. J., Laird, A. R., & Kohn, N. (2025). Emotion regulation: From neural circuits to a transdiagnostic perspective. Neuroscience & Biobehavioral Reviews, 168, 105960. 10.1016/j.neubiorev.2024.105960

Morawetz, C., Riedel, M. C., Salo, T., Berboth, S., Eickhoff, S. B., Laird, A. R., & Kohn, N. (2020). Multiple large-scale neural networks underlying emotion regulation. Neuroscience and Biobehavioral Reviews, 116(July), 382–395. 10.1016/j.neubiorev.2020.07.001

Murphy, K., & Fox, M. D. (2017). Towards a consensus regarding global signal regression for resting state functional connectivity MRI. NeuroImage, 154, 169–173. 10.1016/j.neuroimage.2016.11.052

Ochsner, K. N., Bunge, S. A., Gross, J. J., & Gabrieli, J. D. E. (2002). Rethinking Feelings: An fMRI Study of the Cognitive Regulation of Emotion. Journal of Cognitive Neuroscience, 14(8), 1215–1229. 10.1162/089892902760807212

Ouzzani, M., Hammady, H., Fedorowicz, Z., & Elmagarmid, A. (2016). Rayyan—A web and mobile app for systematic reviews. Systematic Reviews, 5(1), 210. 10.1186/s13643-016-0384-4

Picó-Pérez, M., Radua, J., Steward, T., Menchón, J. M., & Soriano-Mas, C. (2017). Emotion regulation in mood and anxiety disorders: A meta-analysis of fMRI cognitive reappraisal studies. Progress in Neuro-Psychopharmacology and Biological Psychiatry, 79(June), 96–104. 10.1016/j.pnpbp.2017.06.001

Poldrack, R. A. (2011). Inferring Mental States from Neuroimaging Data: From Reverse Inference to Large-Scale Decoding. Neuron, 72(5), 692–697. 10.1016/j.neuron.2011.11.001

Powers, J. P., & LaBar, K. S. (2019). Regulating emotion through distancing: A taxonomy, neurocognitive model, and supporting meta-analysis. Neuroscience and Biobehavioral Reviews, 96(November 2018), 155–173. 10.1016/j.neubiorev.2018.04.023

Preece, D. A., Becerra, R., Robinson, K., & Gross, J. J. (2020). The Emotion Regulation Questionnaire: Psychometric Properties in General Community Samples. Journal of Personality Assessment, 102(3), 348–356. 10.1080/00223891.2018.1564319

Reddan, M. C., Chang, L., Kragel, P., & Wager, T. D. (2024). Somatosensory and motor contributions to emotion representation (arXiv:2411.08973). arXiv. 10.48550/arXiv.2411.08973

Rohrer, J. M., & Murayama, K. (2023). These Are Not the Effects You Are Looking for: Causality and the Within-/Between-Persons Distinction in Longitudinal Data Analysis. Advances in Methods and Practices in Psychological Science, 6(1), 25152459221140842. 10.1177/25152459221140842

Schulz, M.-A., Bzdok, D., Haufe, S., Haynes, J.-D., & Ritter, K. (2024). Performance reserves in brain-imaging-based phenotype prediction. Cell Reports, 43(1). 10.1016/j.celrep.2023.113597

Sheppes, G., Suri, G., & Gross, J. J. (2015). Emotion Regulation and Psychopathology. Annual Review of Clinical Psychology, 11(1), 379–405. 10.1146/annurev-clinpsy-032814-112739

Sicorello, M., Gianaros, P. J., Wright, A. G. C., Petre, B., Kraynak, T., Manuck, S., Schmahl, C., & Wager, T. D. (2025). The functional neurobiology of negative affective traits across regions, networks, signatures, and a machine learning multiverse (p. 2025.05.15.653674). bioRxiv. 10.1101/2025.05.15.653674

Sicorello, M., & Schmahl, C. (2021). Emotion dysregulation in borderline personality disorder: A fronto–limbic imbalance? Current Opinion in Psychology, 37, 114–120. 10.1016/j.copsyc.2020.12.002

Thompson, R. A. (1994). Emotion regulation: A theme in search of definition. Monographs of the Society for Research in Child Development, 59(2–3), 25–52, 250–283. 10.2307/1166137

Vossel, S., Geng, J. J., & Fink, G. R. (2014). Dorsal and Ventral Attention Systems. The Neuroscientist, 20(2), 150–159. 10.1177/1073858413494269

Wager, T. D. (2024). CanlabCore [Computer software]. https://github.com/canlab/CanlabCore

Wager, T. D., Davidson, M. L., Hughes, B. L., Lindquist, M. A., & Ochsner, K. N. (2008). Prefrontal-Subcortical Pathways Mediating Successful Emotion Regulation. Neuron, 59(6), 1037–1050. 10.1016/j.neuron.2008.09.006

Ward, P. G. D., Orchard, E. R., Oldham, S., Arnatkevičiūtė, A., Sforazzini, F., Fornito, A., Storey, E., Egan, G. F., & Jamadar, S. D. (2020). Individual differences in haemoglobin concentration influence bold fMRI functional connectivity and its correlation with cognition. NeuroImage, 221, 117196. 10.1016/j.neuroimage.2020.117196

Zhou, F., Zhao, W., Qi, Z., Geng, Y., Yao, S., Kendrick, K. M., Wager, T. D., & Becker, B. (2021). A distributed fMRI-based signature for the subjective experience of fear. Nature Communications, 12(1), 1–16. 10.1038/s41467-021-26977-3

