## Supplemental Materials for "The elusive neural signature of emotion regulation capabilities: evidence from a large-scale consortium"

### Supplementary Tables

#### Table S1

*Studies included in meta-analytic analyses*

| **Study**<br>First author<br>Publication year | ***N*** | **age**<br>***M* (*SD*)** | **Female (%)** | **Questionnaire:**<br>***M* (*SD*)** | **Stimulus Type** | **Task Ratings** |
|---|---|---|---|---|---|---|
| Benzait et al. (2023) | 17 | 31.4 (9.1) | 35% | ERQ: 4.5 (0.9) | pictures | |
| Berboth et al. (2021) | 25 | 22.8 (3.0) | 84% | ERQ: 2.6 (0.7) | pictures | x |
| Brehl et al. (2021; partially unpublished) | 242 | 22.9 (5.1) | 73% | FEEL-E: 22.7 (4.0) | pictures | x |
| Burghart et al. (unpublished) | 31 | 23.4 (4.1) | 100% | ERQ: 5.0 (0.8) | pictures | x |
| Diers, Gärtner et al. (2023)*; Scheffel et al. (2019) | 39 | 24.6 (4.2) | 66% | ERQ: 4.7 (0.8) | pictures | |
| Dörfel et al. (2014; distancing) | 16 | 23.7 (6.7) | 100% | ERQ: 4.6 (1.1) | pictures | |
| Dörfel et al. (2014; reinterpretation) | 19 | 22.4 (2.7) | 100% | ERQ: 4.4 (1.1) | pictures | |
| Gaebler et al. (2014) | 23 | 30.0 (8.0) | 78% | ERQ: 4.8 (1.2) | pictures | |
| Diers et al. (2021); Gärtner et al. (2019); Scheffel et al. (2019)* | 37 | 25.2 (4.4) | 53% | ERQ: 4.6 (0.9) | pictures | |
| Gianaros et al. (2020; AHAB-II) | 163 | 42.7 (7.1) | 48% | ERQ: 5.0 (0.9) | pictures | x |
| Gianaros et al. (2020; PIP) | 176 | 40.1 (6.3) | 51% | ERQ: 5.0 (0.9) | pictures | x |
| Glosemeyer et al. (2020) | 15 | 24.0 (3.1) | 53% | ERQ: 4.1 (0.9) | cyberball | x |
| Hofhansel et al. (2023) | 16 | 34.7 (10.2) | 0% | NA | game | |
| Jentsch et al. (2019) | 30 | 23.7 (3.4) | 47% | ERQ: 4.5 (1.3) | pictures | x |
| Powers et al. (2022) | 40 | 25.5 (4.7) | 70% | ERQ: 4.8 (0.9) | pictures | x |
| Wessa et al. (unpublished) | 89 | 30.0 (14.0) | 55% | ERQ: 27.0 (6.0) | pictures | x |
| Kim et al. (unpublished) | 33 | 23.0 (2.1) | 52% | ERQ: 4.8 (1.1) | pictures | x |
| LaBar et al. (unpublished) | 53 | 55.0 (12.2) | 62% | ERQ: 4.6 (1.0) | Autobio-graphical memory cues | x |
| Marín-Morales et al. (2022) | 29 | 38.3 (8.2) | 0% | CERQ: 11.9 (11.9) | pictures | x |
| Min et al. (2022) | 105 | 22.8 (2.7) | 49% | ERQ: 29.5 (6.0) | pictures | x |
| Morawetz et al. (2016) | 23 | 22.9 (3.6) | 65% | ERQ: 2.9 (1.1) | videos | x |
| Morawetz et al. (2016; pictures) | 59 | 32.4 (11.2) | 33% | ERQ: 3.0 (1.0) | pictures | x |
| Morawetz et al. (2016; videos) | 59 | 32.4 (11.2) | 33% | ERQ: 3.00 (1.0) | videos | x |

| | | | | | | |
|---|---|---|---|---|---|---|
| Morawetz et al. (2019) | 29 | 24.5 (5.2) | 81% | ERQ: 2.9 (1.1) | pictures | x |
| Morawetz et al. (2020) | 35 | 23.0 (3.4) | 82% | ERQ: 3.1 (1.0) | pictures | x |
| Morawetz et al. (2021) | 37 | 22.0 (3.7) | 87% | ERQ: 2.8 (1.1) | pictures | x |
| Mulej Bratec et al. (2015) | 20 | 24.8 (2.3) | 100% | ERQ: 5.1 (0.5) | pictures | x |
| Müller-Pinzler et al. (unpublished) | 15 | 24.0 (3.12) | 53% | ERQ: 4.0 (0.9) | pictures | x |
| Huneke et al. (unpublished) | 18 | 25.6 (5.6) | 67% | ERQ: 5.4 (0.7) | pictures | x |
| Pierce et al. (2022) | 110 | 43.6 (18.1) | 57% | ERQ: 5.4 (1.2) | pictures | x |
| Paschke et al. (2016) | 115 | 26.1 (3.8) | 51% | ERQ: 3.5 (1.1) | pictures | x |
| Rehbein et al. (2021; sample 1) | 15 | 23.9 (4.6) | 100% | ERQ: 30.8 (3.8) | pictures | x |
| Rehbein et al. (2021; sample 2) | 16 | 23.1 (3.2) | 100% | ERQ: 29.1 (5.3) | pictures | x |
| Sandner et al. (2021) | 38 | 25.0 (4.0) | 50% | CERQ: 15.3 (3.6) | pictures | x |
| Doren et al. (unpublished) | 139 | 39.5 (11.9) | 6% | ERQ: 27.0 (5.5) | pictures | x |
| Guendelman et al. (2022) | 55 | 38.5 (10.0) | 83% | CERQ: 13.9 (3.7) | pictures | x |
| Sokolowski et al. (2022) | 83 | 21.7 (1.8) | 49% | CERQ: 14.6 (3.4) | pictures | x |
| Steward et al. (2021) | 92 | 20.0 (2.8) | 54% | ERQ: 30.5 (6.4) | pictures | x |
| Lloyd et al. (2021) | 59 | 69.5 (7.7) | 50% | CERQ-SF: 6.1 (2.0) | pictures | x |
| Tupitsa et al. (2023) | 19 | 27.0 (5.0) | 55% | CERQ-SF: 8.2 (1.4) | pictures | x |

*Note.* The task ratings column indicates whether task-based affective ratings were present. Only ratings given during the task were considered (e.g., not after the task). Sample size varied slightly depending on the outcome in some studies. Therefore, the sample sizes for questionnaires are reported in this table for simplicity, as these were the lower bound of sample sizes for the three outcomes. For questionnaires authors provided the subscales most indicative of cognitive reappraisal. These were for the reappraisal scale for the ERQ (Emotion Regulation Questionnaire), the positive reappraisal scale for the CERQ (Cognitive Emotion Regulation Questionnaire), and the re-evaluation scale for the FEEL-E (Fragebogen zur Erhebung der Emotionsregulation bei Erwachsenen).
*first-author used for study-identifiers in case the sample was linked to multiple equivalent publications

**Table S2**

*Significant clusters for whole-brain correlations with task-based affective ratings*

| Network | Region | Volume [mm³] | MNI coordinates | | | Max. t-statistic | % covered by region |
|---|---|---|---|---|---|---|---|
| | | | X | Y | Z | | |
| Basal Ganglia | Nucleus Caudatus R | 8 | 22 | 20 | 10 | -3.79 | 100 |
| Brain Stem | Pons L | 104 | -22 | -10 | -8 | -4.88 | 85 |
| Dorsal Attention | Supramarginal Gyrus L | 8 | -64 | -24 | 34 | -3.77 | 100 |
| Dorsal Attention | Supramarginal Gyrus L | 40 | -66 | -26 | 28 | -4.03 | 100 |
| Dorsal Attention | Supramarginal Gyrus L | 48 | -58 | -22 | 22 | -4.13 | 83 |
| Dorsal Attention | Postcentral Gyrus L | 3328 | -44 | -34 | 58 | -7.33 | 31 |
| Somatomotor | Superior Parietal Lobule L | 8 | -22 | -48 | 74 | -3.76 | 100 |
| Somatomotor | Supramarginal Gyrus L | 72 | -62 | -20 | 40 | -4.59 | 56 |
| Somatomotor | Precental Gyrus L | 224 | -28 | -16 | 56 | -4.42 | 32 |
| Somatomotor | Precental Gyrus L | 320 | -30 | -14 | 70 | -4.61 | 73 |
| Somatomotor | Insula Lobe L | 64 | -38 | -4 | 16 | -4.19 | 100 |
| Somatomotor | Rolandic Operculum R | 136 | 56 | -16 | 18 | -4.53 | 94 |
| Somatomotor | Precental Gyrus L | 280 | -58 | 6 | 34 | -4.33 | 74 |
| Visual | Calcarine Gyrus L | 48 | -26 | -62 | 10 | -3.96 | 33 |

*Note.* Labelling was performed using the automated CanlabCore atlas-based procedure combined with the SPM12 anatomy toolbox after exclusion of Brehl et al. (2021).

### Supplementary Figures

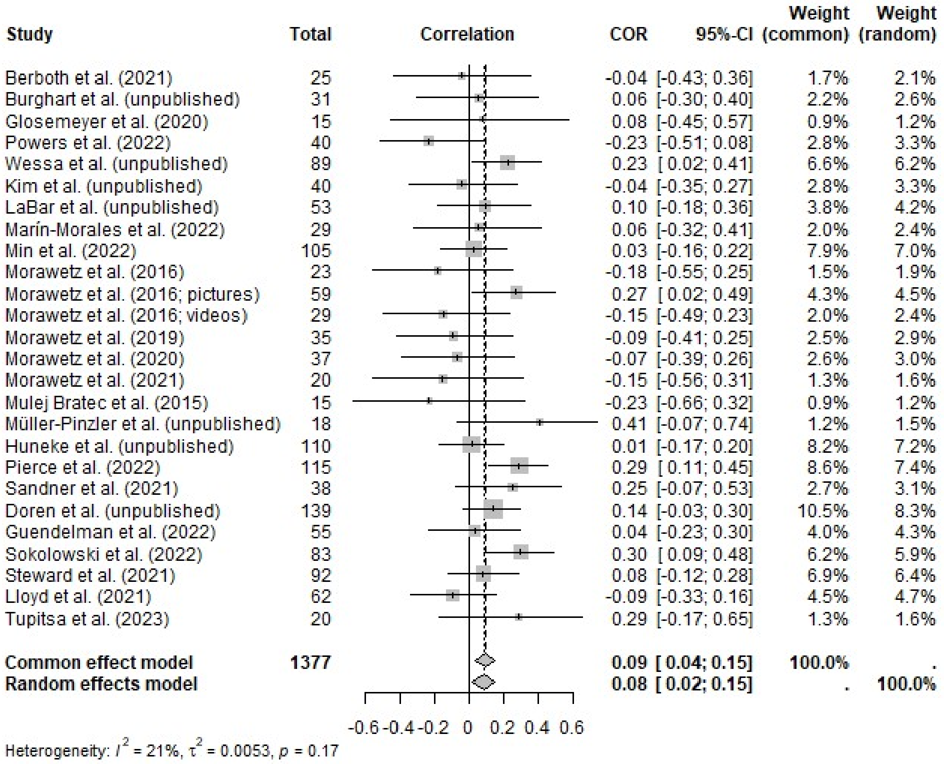

**Figure S1.**

Forest plot of correlations between amygdala down-regulation and task-based affective ratings.

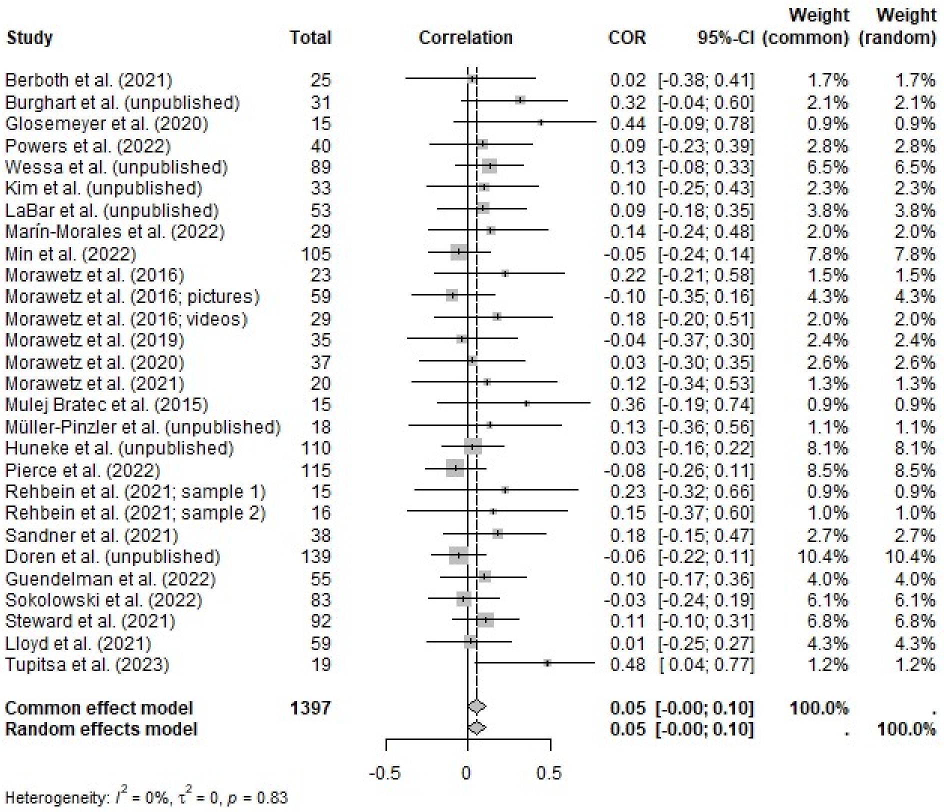

**Figure S2.**

Forest plot of correlations between trait questionnaires and task-based affective ratings.

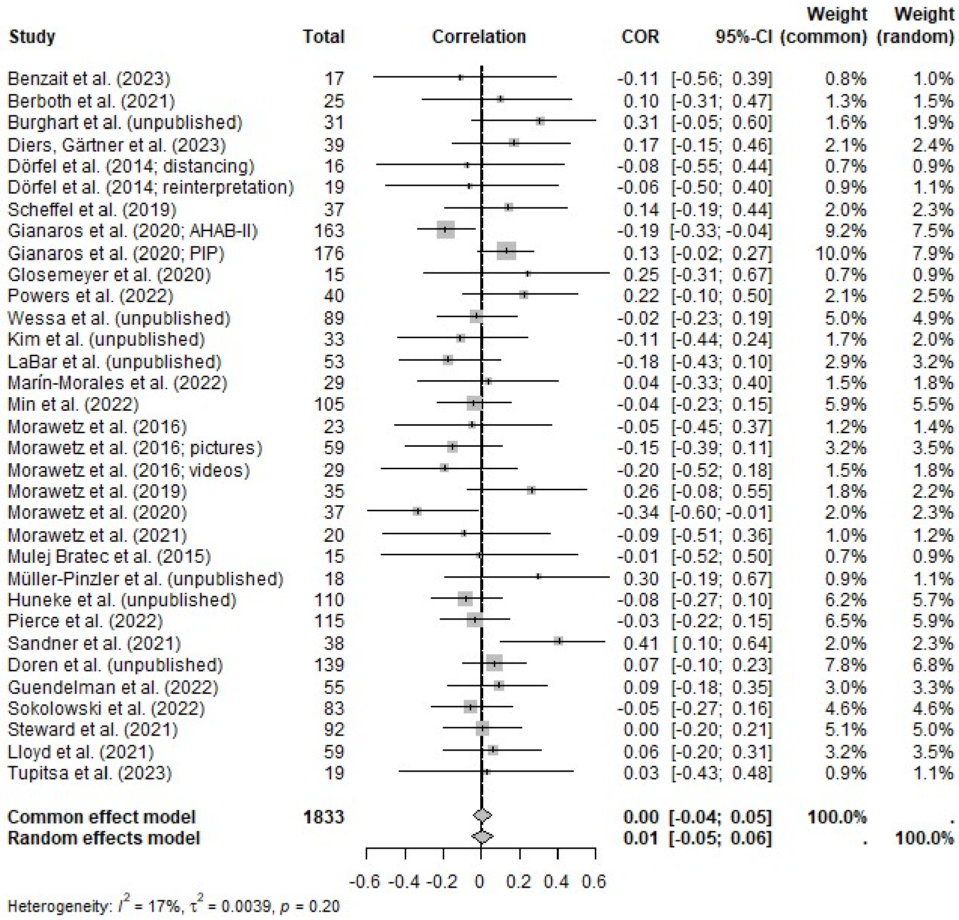

**Figure S3.**

Forest plot of correlations between trait questionnaires and amygdala down-regulation.

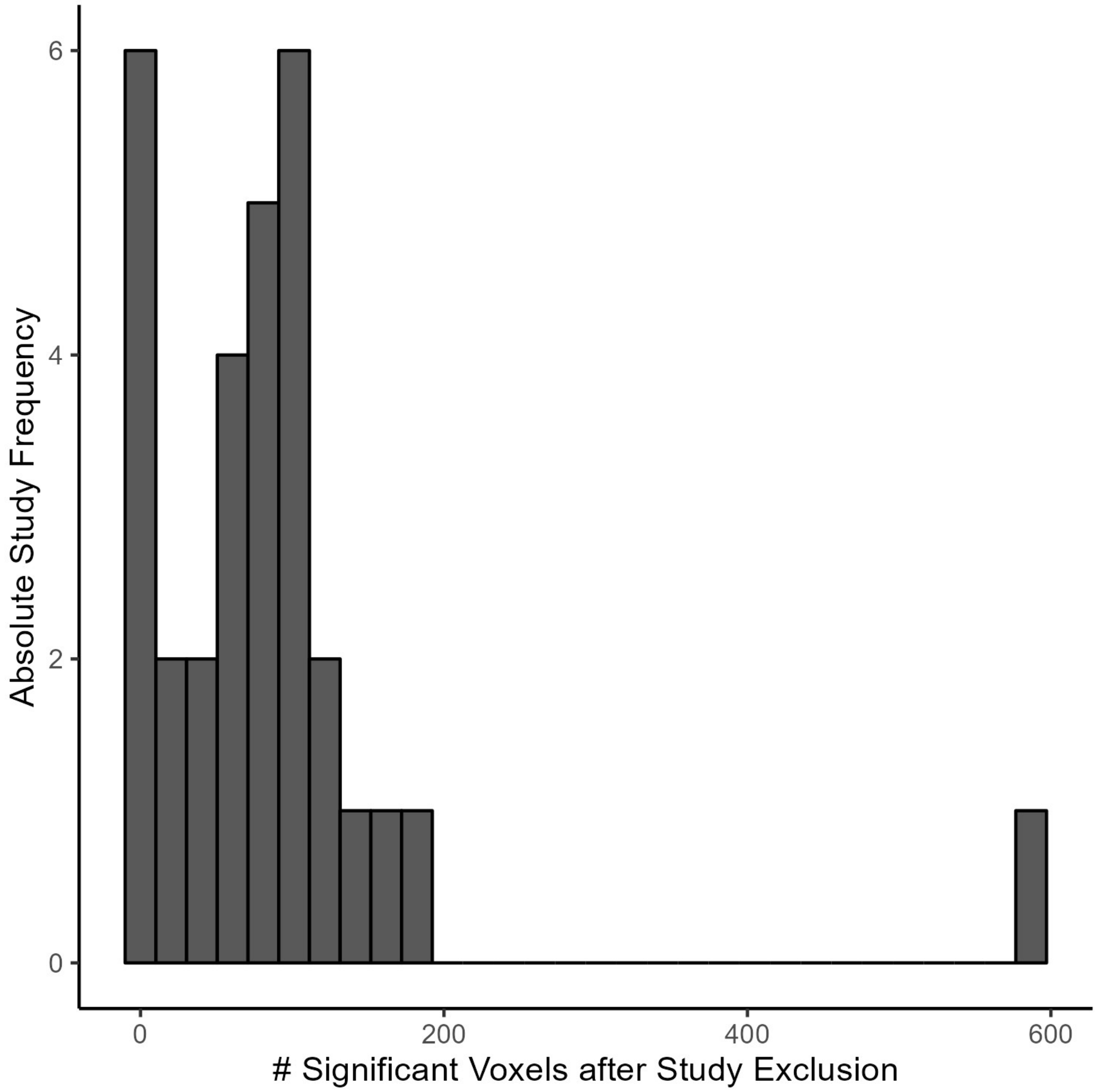

**Figure S4.**

Results of the jackknife procedure for task-based affective ratings. Changes in number of significant voxels after excluding a single-study. The column on the far right only consists of the Brehl et al. (2021) study.

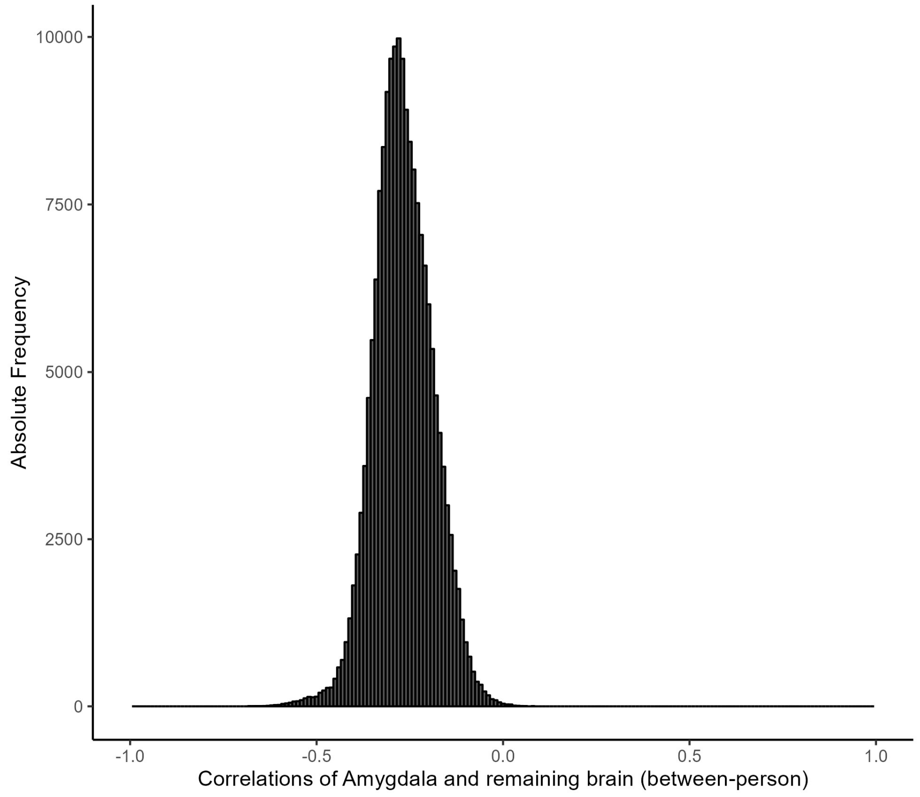

**Figure S5.**

Between-person correlations of amygdala activation in the contrast [view- regulate] and the rest of the brain in the reverse contrast [regulate - view]. For simplicity, the main paper reports the correlations in the same contrast [regulate - view].

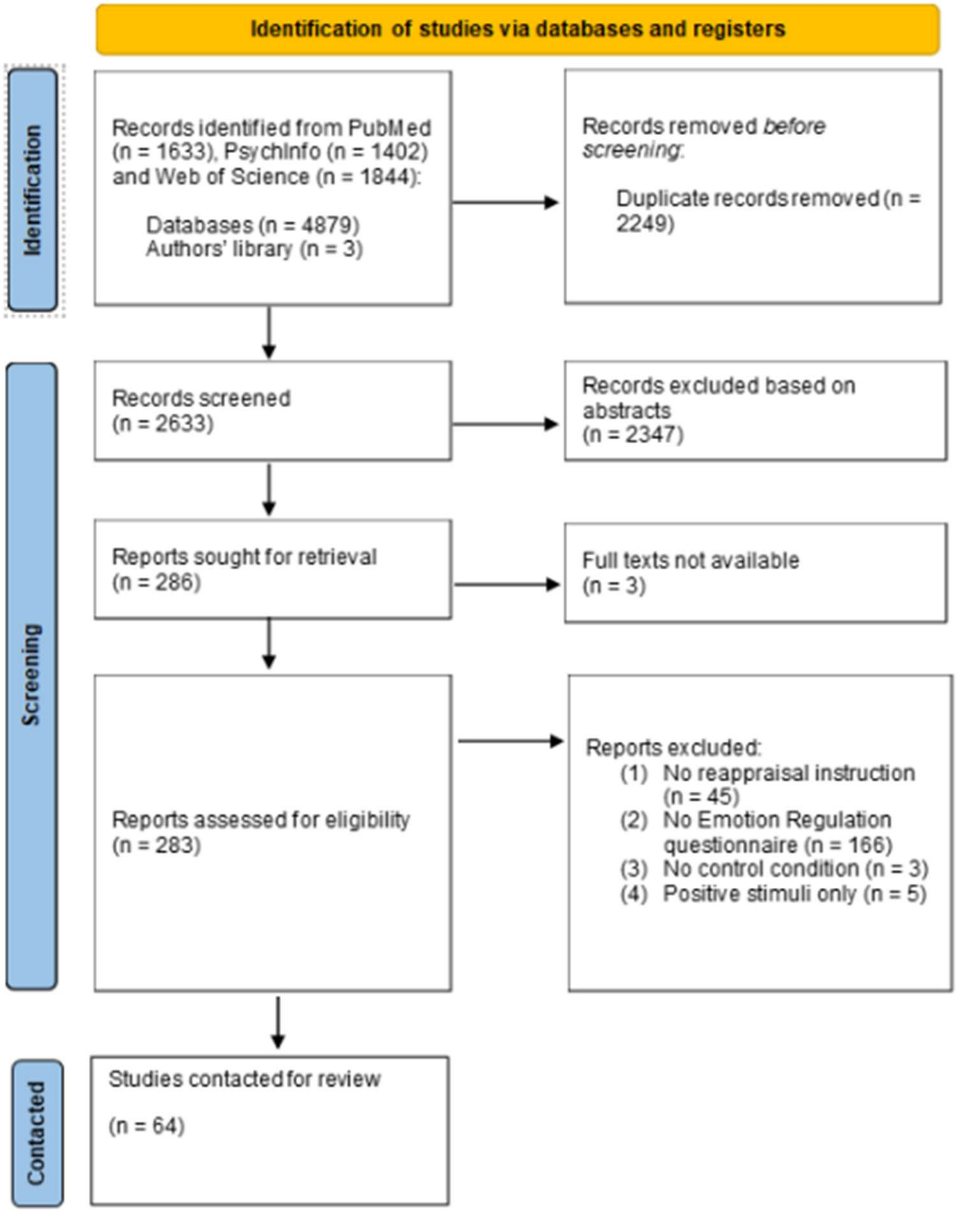

**Figure S6.**

Flowchart of literature search and screening procedure.

### Supplementary Methods

#### Testing the custom meta-analytic functions

All steps for code quality checks can be found and reproduced using the matlab file "testMetaAnalysisFunction.m" on the github repository. The meta-analytic code was tested in three steps. First, we tested the convergence of the core function with the results from the established R package *meta* and an in-built dataset, indicating perfect agreement (Balduzzi et al., 2019). Second, we assigned this dataset to an fmri_data object and tested whether our image-based function leads to identical results. Here, a deviation after seven decimal places was observed, which was due to the conversion from double to single precision arrays when using canlabCore fmri_data objects. This is not a problem when using fMRI-based image data, which are loaded with single precision by default. Third, we searched for openly accessible study-wise group-level t-maps on neurovault for the contrast [painful stimulus - non-painful stimulus] to perform a small ad-hoc meta-analysis. We chose pain, as its neurobiology is relatively well known and core regions replicate relatively well across different designs, making it a viable ground truth model. Here, we found seven matching images. One was removed due to insufficient coverage, leading to a sample size of $k = 6$ and $N = 297$. A list of permanent neurovault identifiers to retrieve the images can be found in the file "studyInfo_neurovault.xlsx" in the github repository. The results of the final code for image-based random effects meta-analysis led to converging results with a recently published image-based meta-analysis on pain (Figure S7; Zunhammer et al., 2021) and was decoded as *pain* using neurosynth at a image-topic correlation of $r = .44$, which is a relatively large correlation in our experience with this procedure. The results from Zunhammer et al. (2021) were slightly above this decoding precision ($r = .54$), which should be expected based on their larger sample size.

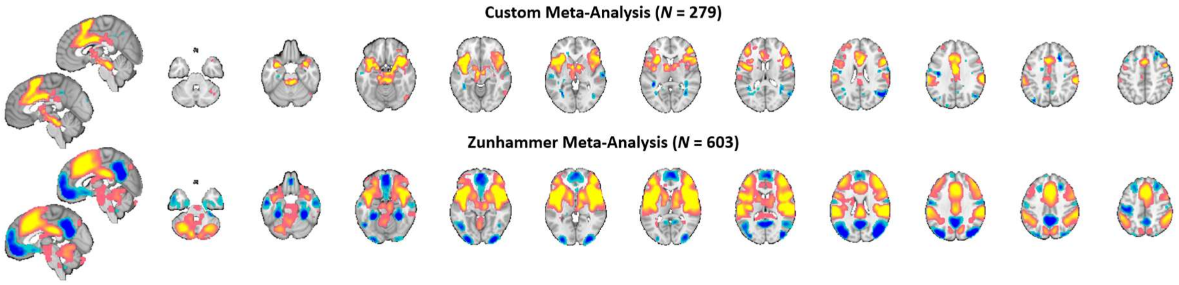

**Figure S7.** Comparison of a small ad-hoc meta-analysis with our custom code and a meta-analysis by Zunhammer and colleagues (2021). Note that the latter had a much larger sample size and was based on a systematic literature review, explaining the larger number of significant effects.

**Power Analysis**

We performed detailed power simulations to assess which range of hypothetical effect sizes our sample can detect. We built custom functions which estimate the power for detecting at least one true positive voxel and takes as input a target effect size, a vector of study-wise sample sizes, the between-study heterogeneity [tau], a statistical threshold, and the number of voxels hypothesized to have an effect of the specified size.

Our procedure followed a four-step logic. First, the significance threshold for at least one voxel surviving FDR correction is 0.05/V, where V is the number of voxels tested (e.g., ≈10,000 in our network of interest and ≈180,000 in the whole brain). Secondly, analytic formulas for statistical power in random effects meta-analyses can be easily implemented for single tests using this corrected threshold (Jackson & Turner, 2017). Thirdly, the more voxels have a true correlation in the population, the higher the chance that at least one true-positive voxel will cross the threshold. If voxels were independent tests, the probability for this "family-wise power" (FWP) is FWP = $1-\beta^k$, where $\beta$ is the false negative rate for a single voxel and k is the number of voxels assumed to have an effect. Still, as voxels are usually highly dependent, FWP will be strongly overestimated (a stochastic phenomenon usually described in the context of false positive rates; Nichols & Hayasaka, 2003). Therefore, fourthly, we estimated the effective number of tests and replaced this number for k. More precisely, we sampled different numbers of voxels from our network of interest (10, 100, 1000, 10000), computed a correlation matrix for these voxels from our dataset of brain-questionnaire correlations, and estimated the effective number of tests using the procedure by Li and Ji (2005), which was found to be the most accurate approach for high-dimensional genetic data (Wen & Lu, 2011).

We tested this procedure by simulating 10,000 datasets with $N = 250$ from an empirical correlation matrix of 20 voxels from our network of interest in addition to a synthetic outcome variable correlated at r = 0.1 with all 20 voxels. The estimated ground truth power was 58% and slightly overestimated with our procedure at 64%, potentially implying slightly positively biased results.

For each iteration, we calculated the effective number of tests using four different methods and, using this estimate, the hypothetical power for the predefined effect size (r = 0.10) with FDR correction for 20 tests. These power estimates were averaged over iterations. As ground truth, we calculated for each iteration whether at least one "voxel" was significantly correlated to the outcome variable after FDR correction and calculated power as the number of iterations with at least one positive test after correction divided by the total number of iterations. We found that all four methods slightly overestimated power, with the smallest difference between the Galway method (62%) and ground truth (58%), followed by the Li method (64%). We still chose the Li method, due to concordance with previous findings and the observation that the number of effective tests had a ceiling at already 100 voxels for the Galway method. Our simulation procedure could not test the accuracies of the procedures for these numbers of voxels, due to multicollinearity in empirical correlation matrices above 20 voxels.

### References

Balduzzi, S., Rücker, G., & Schwarzer, G. (2019). How to perform a meta-analysis with R: A practical tutorial. *Evidence-Based Mental Health*, *22*(4), 153–160. https://doi.org/10.1136/ebmental-2019-300117

Benzait, A., Krenz, V., Wegrzyn, M., Doll, A., Woermann, F., Labudda, K., Bien, C. G., & Kissler, J. (2023). Hemodynamic correlates of emotion regulation in frontal lobe epilepsy patients and healthy participants. *Human Brain Mapping*, *44*(4), 1456–1475. https://doi.org/10.1002/hbm.26133

Berboth, S., Windischberger, C., Kohn, N., & Morawetz, C. (2021). Test-retest reliability of emotion regulation networks using fMRI at ultra-high magnetic field. *NeuroImage*, *232*, 117917. https://doi.org/10.1016/j.neuroimage.2021.117917

Brehl, A.-K., Schene, A., Kohn, N., & Fernández, G. (2021). Maladaptive emotion regulation strategies in a vulnerable population predict increased anxiety during the Covid-19 pandemic: A pseudo-prospective study. *Journal of Affective Disorders Reports*, *4*, 100113. https://doi.org/10.1016/j.jadr.2021.100113

Diers, K.*, Gärtner, A.*, Schönfeld, S., Dörfel, D., Walter, H., Brocke, B., & Strobel, A. (2023). Short- and long-term effects of emotion up- and down-regulation. *Imaging Neuroscience*, *1*, 1–24. https://doi.org/10.1162/imag_a_00028

Diers, K., Dörfel, D., Gärtner, A., Schönfeld, S., Walter, H., Strobel, A., & Brocke, B. (2021). Should we keep some distance from distancing? Regulatory and post-regulatory effects of emotion downregulation. *PLOS ONE, 16*(9), e0255800. https://doi.org/10.1371/journal.pone.0255800

Dörfel, D., Lamke, J.-P., Hummel, F., Wagner, U., Erk, S., & Walter, H. (2014). Common and differential neural networks of emotion regulation by Detachment, Reinterpretation, Distraction, and Expressive Suppression: A comparative fMRI investigation. *NeuroImage*, *101*, 298–309. https://doi.org/10.1016/j.neuroimage.2014.06.051

Gärtner, A., Dörfel, D., Diers, K., Witt, S. H., Strobel, A., & Brocke, B. (2019). Impact of FAAH genetic variation on fronto-amygdala function during emotional processing. *European Archives of Psychiatry and Clinical Neuroscience, 269*(2), 209–221. https://doi.org/10.1007/s00406-018-0944-9

Gianaros, P. J., Kraynak, T. E., Kuan, D. C.-H., Gross, J. J., McRae, K., Hariri, A. R., Manuck, S. B., Rasero, J., & Verstynen, T. D. (2020). Affective brain patterns as multivariate neural correlates of cardiovascular disease risk. *Social Cognitive and Affective Neuroscience*, *March*, 1–12. https://doi.org/10.1093/scan/nsaa050

Glosemeyer, R. W., Diekelmann, S., Cassel, W., Kesper, K., Koehler, U., Westermann, S., Steffen, A., Borgwardt, S., Wilhelm, I., Müller-Pinzler, L., Paulus, F. M., Krach, S., & Stolz, D. S. (2020). Selective suppression of rapid eye movement sleep increases next-day negative affect and amygdala responses to social exclusion. *Scientific Reports*, *10*(1), 17325. https://doi.org/10.1038/s41598-020-74169-8

Guendelman, S., Bayer, M., Prehn, K., & Dziobek, I. (2022). Towards a mechanistic understanding of mindfulness-based stress reduction (MBSR) using an RCT neuroimaging approach: Effects on regulating own stress in social and non-social situations. *NeuroImage*, *254*, 119059. https://doi.org/10.1016/j.neuroimage.2022.119059

Hofhansel, L., Weidler, C., Clemens, B., Habel, U., & Votinov, M. (2023). Personal insult disrupts regulatory brain networks in violent offenders. *Cerebral Cortex*, *33*(8), 4654–4664. https://doi.org/10.1093/cercor/bhac369

Jackson, D., & Turner, R. (2017). Power analysis for random-effects meta-analysis. *Research Synthesis Methods*, *8*(3), 290–302. https://doi.org/10.1002/jrsm.1240

Jentsch, V. L., Merz, C. J., & Wolf, O. T. (2019). Restoring emotional stability: Cortisol effects on the neural network of cognitive emotion regulation. *Behavioural Brain Research*, *374*, 111880. https://doi.org/10.1016/j.bbr.2019.03.049

Li, J., & Ji, L. (2005). Adjusting multiple testing in multilocus analyses using the eigenvalues of a correlation matrix. *Heredity*, *95*(3), 221–227. https://doi.org/10.1038/sj.hdy.6800717

Lloyd, W. K., Morriss, J., Macdonald, B., Joanknecht, K., Nihouarn, J., & van Reekum, C. M. (2021). Longitudinal change in executive function is associated with impaired top-down frontolimbic regulation during reappraisal in older adults. *NeuroImage*, *225*, 117488. https://doi.org/10.1016/j.neuroimage.2020.117488

Marín-Morales, A., Pérez-García, M., Catena-Martínez, A., & Verdejo-Román, J. (2022). Emotional Regulation in Male Batterers When Faced With Pictures of Intimate Partner Violence. Do They Have a Problem With Suppressing or Experiencing Emotions? *Journal of Interpersonal Violence*, *37*(11–12), NP10271–NP10295. https://doi.org/10.1177/0886260520985484

Min, J., Nashiro, K., Yoo, H. J., Cho, C., Nasseri, P., Bachman, S. L., Porat, S., Thayer, J. F., Chang, C., Lee, T.-H., & Mather, M. (2022). Emotion Downregulation Targets Interoceptive Brain Regions While Emotion Upregulation Targets Other Affective Brain Regions. *Journal of Neuroscience*, *42*(14), 2973–2985. https://doi.org/10.1523/JNEUROSCI.1865-21.2022

Morawetz, C., Berboth, S., & Bode, S. (2021). With a little help from my friends: The effect of social proximity on emotion regulation-related brain activity. *NeuroImage*, *230*, 117817. https://doi.org/10.1016/j.neuroimage.2021.117817

Morawetz, C., Bode, S., Baudewig, J., Jacobs, A. M., & Heekeren, H. R. (2016). Neural representation of emotion regulation goals. *Human Brain Mapping*, *37*(2), 600–620.

https://doi.org/10.1002/hbm.23053

Morawetz, C., Bode, S., Baudewig, J., Kirilina, E., & Heekeren, H. R. (2016). Changes in Effective Connectivity Between Dorsal and Ventral Prefrontal Regions Moderate Emotion Regulation. *Cerebral Cortex*, *26*(5), 1923–1937. https://doi.org/10.1093/cercor/bhv005

Morawetz, C., Mohr, P. N. C., Heekeren, H. R., & Bode, S. (2019). The effect of emotion regulation on risk-taking and decision-related activity in prefrontal cortex. *Social Cognitive and Affective Neuroscience*, *14*(10), 1109. https://doi.org/10.1093/scan/nsz078

Morawetz, C., Steyrl, D., Berboth, S., Heekeren, H. R., & Bode, S. (2020). Emotion Regulation Modulates Dietary Decision-Making via Activity in the Prefrontal–Striatal Valuation System. *Cerebral Cortex (New York, NY)*, *30*(11), 5731. https://doi.org/10.1093/cercor/bhaa147

Mulej Bratec, S., Xie, X., Schmid, G., Doll, A., Schilbach, L., Zimmer, C., Wohlschläger, A., Riedl, V., & Sorg, C. (2015). Cognitive emotion regulation enhances aversive prediction error activity while reducing emotional responses. *NeuroImage*, *123*, 138–148. https://doi.org/10.1016/j.neuroimage.2015.08.038

Nichols, T., & Hayasaka, S. (2003). Controlling the familywise error rate in functional neuroimaging: A comparative review. *Statistical Methods in Medical Research*, *12*(5), 419–446. https://doi.org/10.1191/0962280203sm341ra

Paschke, L. M., Dörfel, D., Steimke, R., Trempler, I., Magrabi, A., Ludwig, V. U., Schubert, T., Stelzel, C., & Walter, H. (2016). Individual differences in self-reported self-control predict successful emotion regulation. *Social Cognitive and Affective Neuroscience*, *11*(8), 1193–1204. https://doi.org/10.1093/scan/nsw036

Pierce, J. E., Blair, R. J. R., Clark, K. R., & Neta, M. (2022). Reappraisal-related downregulation of amygdala BOLD activation occurs only during the late trial window. *Cognitive, Affective, & Behavioral Neuroscience*, *22*(4), 777–787. https://doi.org/10.3758/s13415-021-00980-z

Powers, J. P., Kako, N., McIntosh, D. N., & McRae, K. (2022). Competitive interactions between cognitive reappraisal and mentalizing. *International Journal of Psychophysiology*, *174*, 17–28. https://doi.org/10.1016/j.ijpsycho.2022.01.012

Rehbein, E., Kogler, L., Hornung, J., Morawetz, C., Bayer, J., Krylova, M., Sundström-Poromaa, I., & Derntl, B. (2021). Estradiol administration modulates neural emotion regulation. *Psychoneuroendocrinology*, *134*, 105425. https://doi.org/10.1016/j.psyneuen.2021.105425

Sandner, M., Zeier, P., Lois, G., & Wessa, M. (2021). Cognitive emotion regulation withstands the stress test: An fMRI study on the effect of acute stress on distraction and reappraisal. *Neuropsychologia*, *157*, 107876. https://doi.org/10.1016/j.neuropsychologia.2021.107876

Scheffel, C., Diers, K., Schönfeld, S., Brocke, B., Strobel, A., & Dörfel, D. (2019). Cognitive emotion regulation and personality: An analysis of individual differences in the neural and behavioral correlates of successful reappraisal. *Personality Neuroscience*, *2*. https://doi.org/10.1017/pen.2019.11

Sokołowski, A., Morawetz, C., Folkierska-Żukowska, M., & Łukasz Dragan, W. (2022). Brain activation during cognitive reappraisal depending on regulation goals and stimulus valence. *Social Cognitive and Affective Neuroscience*, *17*(6), 559-570.

Steward, T., Davey, C. G., Jamieson, A. J., Stephanou, K., Soriano-Mas, C., Felmingham, K. L., & Harrison, B. J. (2021). Dynamic Neural Interactions Supporting the Cognitive Reappraisal of Emotion. *Cerebral Cortex*, *31*(2), 961–973. https://doi.org/10.1093/cercor/bhaa268

Tupitsa, E., Egbuniwe, I., Lloyd, W. K., Puertollano, M., Macdonald, B., Joanknecht, K., Sakaki, M., & van Reekum, C. M. (2023). Heart rate variability covaries with amygdala functional connectivity during voluntary emotion regulation. *NeuroImage*, *274*, 120136. https://doi.org/10.1016/j.neuroimage.2023.120136

Wen, S.-H., & Lu, Z.-S. (2011). Factors affecting the effective number of tests in genetic association studies: A comparative study of three PCA-based methods. *Journal of Human Genetics*, *56*(6), 428–435. https://doi.org/10.1038/jhg.2011.34

Zunhammer, M., Spisák, T., Wager, T. D., Bingel, U., & Placebo Imaging Consortium. (2021). Meta-analysis of neural systems underlying placebo analgesia from individual participant fMRI data. *Nature Communications*, *12*(1), 1391. https://doi.org/10.1038/s41467-021-21179-3
